# Innate immune signaling in *Drosophila* shifts anabolic lipid metabolism from triglyceride storage to phospholipid synthesis in an ER stress-dependent manner

**DOI:** 10.1101/2020.03.01.972208

**Authors:** Brittany A. Martínez, Scott Yeudall, Rosalie G. Hoyle, J. David Castle, Norbert Leitinger, Michelle L. Bland

## Abstract

During infection, cellular resources are allocated toward the metabolically-demanding processes of synthesizing and secreting effector proteins that neutralize and kill invading pathogens. In *Drosophila*, these effectors are antimicrobial peptides (AMPs) that are produced in the fat body, an organ that also serves as a major lipid storage depot. Here we asked how activation of Toll signaling in the larval fat body perturbs lipid homeostasis to understand how cells meet the metabolic demands of the immune response. We find that genetic activation of fat body Toll signaling leads to a tissue-autonomous reduction in triglyceride storage that is paralleled by decreased transcript levels of *Lipin*, which synthesizes diacylglycerol, and *midway*, which carries out the final step of triglyceride synthesis. In contrast, we discovered that Kennedy pathway enzymes, such as easily shocked and Pcyt1, that synthesize membrane phospholipids are induced by the Toll pathway. Mass spectrometry analysis revealed elevated levels of major phosphatidylcholine and phosphatidylethanolamine species in fat bodies with active Toll signaling. The induction of Kennedy pathway enzymes in response to Toll signaling required the unfolded response mediator Xbp1 but was blunted by deleting AMP genes and thereby reducing secretory demand elicited by Toll activation. Consistent with these findings, endoplasmic reticulum volume is expanded in fat body cells with active Toll signaling, as determined by transmission electron microscopy. Our results establish that Toll signaling induces a shift in anabolic lipid metabolism, accompanied by changes in key lipid synthesis enzymes, that may serve the immediate demand for AMP synthesis and secretion but that ultimately leads to the long-term consequence of insufficient nutrient storage.

**Author summary:** Fighting infection requires that immune cells synthesize antimicrobial peptides and antibodies and carry out cellular processes like phagocytosis to destroy microbes and clear infected cells. During infection, metabolic processes support and direct immune function. Here, we use the fruit fly Drosophila melanogaster as a model system to understand the interaction between immunity and metabolism. In Drosophila larvae, infection leads to tremendous production of antimicrobial peptides that destroy invading microbes. These peptides are made in the fat body, an organ that is also the site of fat storage. Activating the immune response reduces lipid storage but increases the production of phospholipids that form the membranes of organelles such as the endoplasmic reticulum. This organelle is the starting point for synthesis and secretion of antimicrobial peptides, and its volume is increased in response to immune activation. Shifting metabolism from fat storage to membrane phospholipid synthesis supports the immune response. However, this comes at the expense of the ability to withstand other types of stress such as food scarcity. These findings are important because they suggest that some of the metabolic changes induced by fighting infection may become pathological if they are maintained over long periods of time.

## Introduction

Animals fight viral, microbial and parasitic infections by activating humoral and cellular immune processes. Work in invertebrate and mammalian systems over many decades has identified the molecular pathways that lead from pathogen recognition to synthesis of effector proteins and activation of cellular processes that destroy microbes and infected cells. For example, presentation of pathogen-expressed molecules to conserved host pattern recognition receptors, such as *Drosophila* Toll and mammalian Toll-like receptors (TLRs), activates NF-κB family transcription factors that direct expression of antimicrobial peptides, acute phase proteins and cytokines. These effector proteins rupture microbial membranes, participate in opsonization, lysis and clotting reactions and promote inflammation and activation of other immune cell types. In animals with adaptive immune systems, lymphocytes recognize pathogens via expression of recombinant receptors leading to clonal expansion, antibody secretion and killing of infected cells.

Profound changes in host metabolism accompany the synthesis of antimicrobial peptides, acute phase proteins, cytokines, and antibodies as well as the induction of cellular processes such as phagocytosis and immune cell proliferation during the immune response. These metabolic changes support tolerance of and resistance to infection. At the whole-animal level, for example, rodents injected with lipopolysaccharide (LPS) or infected with *Escherichia coli* exhibit reductions in core temperature and oxygen consumption that drive disease tolerance [1,2]. At the level of individual cells, immune signaling shifts glucose metabolism from oxidative phosphorylation to aerobic glycolysis; a switch that promotes cytokine synthesis as well as survival, activation and bactericidal functions of immune cell types in flies and mammals [3-6]. In dendritic cells stimulated with LPS, aerobic glycolysis drives fatty acid synthesis that underlies expansion of the endoplasmic reticulum (ER) [7]. Changes in lipid metabolism promote secretory function in multiple immune cell types. Upon simulation with LPS, mouse B cells differentiate into plasma cells that synthesize and secrete antibodies; this is accompanied by increased membrane phospholipid synthesis and ER expansion [8,9]. These changes are dependent on splicing of the *X-box binding protein 1* (*Xbp1*) mRNA, leading to a mature transcript that encodes the Xbp1 transcription factor, a key mediator of the unfolded protein response [10,11]. Similarly, in macrophages, phosphatidylcholine synthesis and Xbp1 activation are necessary for maximal levels of cytokine secretion in response to infection or LPS stimulation [12,13]. The mechanisms linking immune signaling with regulation of carbohydrate and lipid metabolic pathways, whether in the physiological response to infection or during pathological disease states characterized by chronic inflammation remain unclear in many cases.

Fighting infection is energetically demanding, and, as in mammals, activation of innate immune signaling in *Drosophila* alters metabolism. Flies respond to septic injury with Gram-positive bacteria or fungi by activating a conserved Toll-nuclear factor-κB (NF-κB) pathway that stimulates synthesis and secretion of micromolar quantities of antimicrobial peptides (AMPs) into hemolymph [14,15]. AMPs such as Drosomycin fight infection by disrupting microbial membranes and inhibiting fungal spore germination [16,17]. The major source of circulating AMPs in infected *Drosophila* larvae is the fat body, an organ that coordinates not only the humoral immune response via activation of the Toll and Imd signaling pathways but also nutrient storage and animal growth [14,18]. Metabolism and growth are regulated by *Drosophila* insulin-like peptides that bind to insulin receptors on fat body cells, leading to activation of Akt and mTOR to stimulate protein synthesis, cell growth, and storage of dietary sugar as triglycerides and glycogen via highly-conserved metabolic pathways [19]. The coordination of immune, growth and metabolic pathways in the same cells of the fat body and the sequence and functional conservation of these pathways between flies and mammals make *Drosophila* an attractive model to study immunometabolism at the scale of whole-animal physiology. Infection of adult flies with the intracellular bacterial pathogen *Mycobacterium marinarum* leads to a progressive depletion of whole-animal triglyceride stores concomitant with lipid accumulation in phagocytes that harbor mycobacteria [20,21]. Colonization of adult fat body cells with the intracellular parasite *Tubulinosema ratisbonensis* also impairs triglyceride storage, directing host fatty acids to fuel parasite growth [22]. Lipid storage defects can be elicited by genetic activation of the Toll and Imd pathways, indicating that metabolic changes are dictated not only by pathogen interaction but also by signaling from the host immune system. Activation of the Imd pathway in larval fat body results in decreased triglyceride levels and impaired whole-animal growth (Davoodi et al., 2019). Expression of a constitutively-active Toll receptor, Toll^10b^, in larval fat body inhibits whole-animal growth, disrupts insulin signaling in fat body, and reduces triglyceride storage [23-25].

While the signaling events that lead from pathogen recognition to immune effector production are well understood, the mechanisms that underlie altered lipid metabolism in response to immune signaling and the short- and long-term consequences of such metabolic changes remain unclear. To address this, we genetically activated Toll signaling in the larval fat body and assessed expression of enzymes that carry out *de novo* lipogenesis. We find that the decrease in triglycerides caused by chronic Toll signaling is mirrored by a selective decrease in expression of enzymes that carry out the two final steps of *de novo* triglyceride synthesis: the phosphatidic acid phosphatase Lipin and the diacylglycerol transferase homolog midway. Enzymes that carry out early steps of fatty acid synthesis are unchanged in response to Toll signaling, leading us to investigate other fates of fatty acids. We find that Toll signaling induces a coordinated increase in expression of enzymes in the Kennedy phospholipid synthesis pathway and elevated levels of the membrane phospholipids phosphatidylethanolamine and phosphatidylcholine. Induction of Kennedy pathway enzymes by Toll receptor activation is dependent on the transcription factor Xbp1, suggesting a contribution of ER stress to this phenotype. Indeed, the ER is expanded and dilated in fat body cells with active Toll signaling. Furthermore, loss of genes encoding AMPs blunts induction of Kennedy pathway enzymes in immune-activated fat body cells. Our results suggest a mechanism by which phospholipid biosynthesis, induced downstream of Toll receptor activation, supports ER expansion to sustain secretory capacity during the immune response. Our data also indicate that a long-term consequence of this metabolic switch that directs fatty acids toward phospholipid synthesis is reduced nutrient storage.

## Results

### Toll signaling in fat body acts in a tissue-autonomous manner to disrupt nutrient storage

The fat body stores the bulk of triglycerides in fruit fly larvae. Activating Toll signaling in larval fat body, via r4-GAL4 driven expression of constitutively-active Toll^10b^, decreased late third instar fat body triglyceride levels by 55%, but led to negligible changes in gut and carcass triglycerides compared with tissues from control larvae expressing GFP in fat body (Fig 1A). We also note, as expected, that the fat body stored 60-80 times more triglyceride than the gut and the carcass, which comprises cuticle, muscle, oenocytes, imaginal discs and trachea. Throughout the larval third instar, animals with active fat body Toll signaling consistently stored less triglyceride than GFP-expressing controls. This held true whether triglyceride levels were normalized to whole-animal protein levels (Fig 1B) or body weight (S1A Fig). Whole-animal protein levels were not altered by fat body Toll signaling over the same time course (S1B Fig). Reduced triglyceride storage in animals with active Toll signaling persisted to the white prepupal stage, a well-defined developmental endpoint that follows the cessation of feeding, indicating that low triglyceride levels are not due to a developmental delay. To understand whether the decrease in triglycerides elicited by innate immune signaling is transcriptionally regulated, we manipulated the NF-κB homolog Dif. Dif is activated by Toll signaling and contributes to the negative regulation of whole-animal growth downstream of Toll activation [25]. Elevated expression of Dif in larval fat body phenocopied Toll^10b^, leading to a decrease in whole-animal triglyceride levels (Fig 1C). However, loss of Dif in fat bodies with active Toll signaling rescued triglyceride storage (Fig 1D).

**Fig 1.**
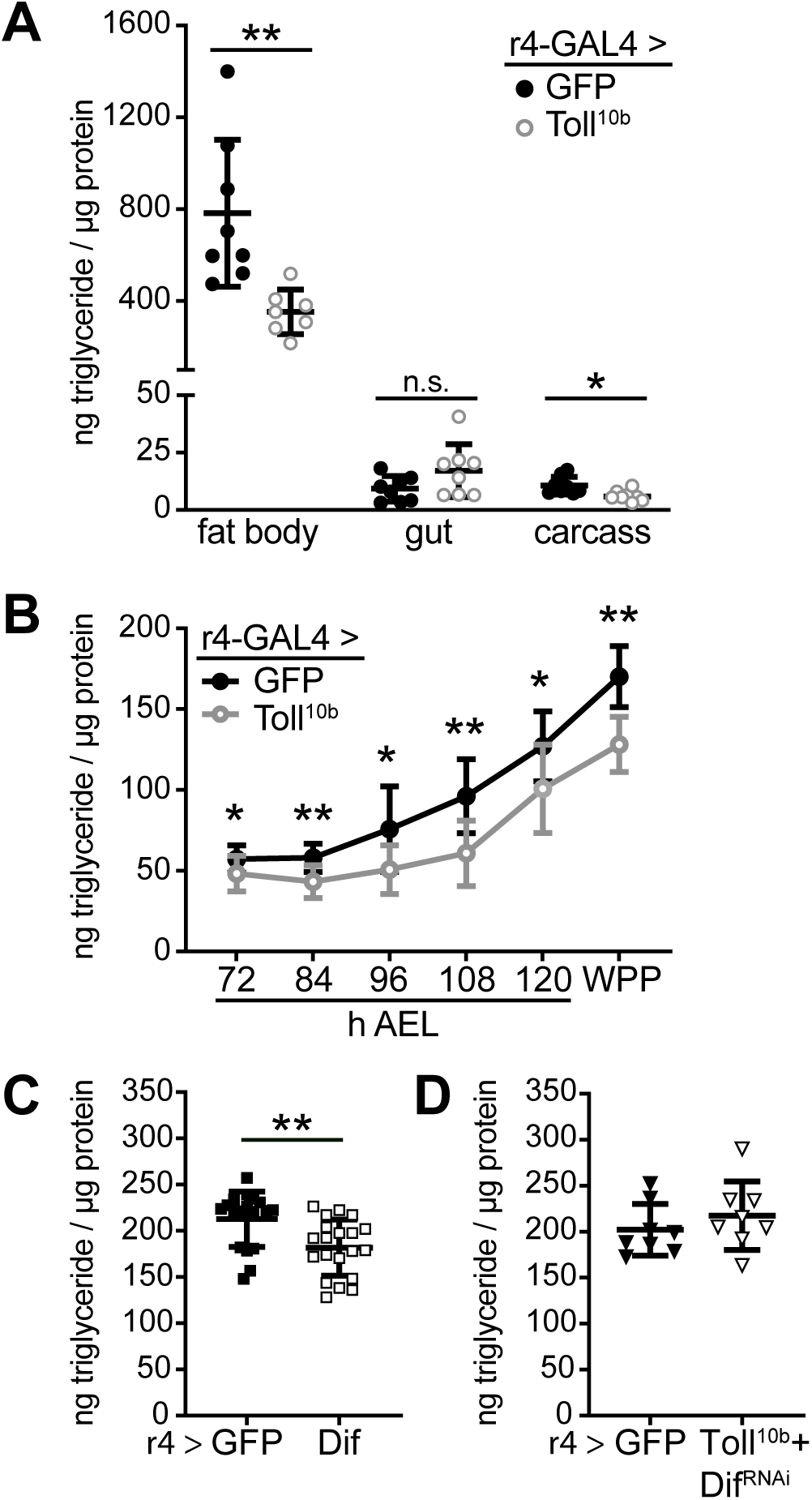
Toll signaling in the third instar larval fat body reduces triglyceride storage in a tissue-autonomous manner. r4-GAL4 was used to drive indicated transgenes in fat body, and triglyceride levels were measured in whole larvae or dissected organs and normalized to protein levels. (**A**) Triglyceride levels in fat body, gut, or carcass, n = 7-8/group. *p = 0.0118 and **p = 0.0047 versus GFP. (**B**) Whole-animal triglyceride levels throughout the third instar (72-120 hours after egg lay (h AEL)) and in white prepupae (WPP), n = 10-11/group. *p ≤ 0.0459 and **p ≤ 0.0019 versus GFP. (**C**) Whole-animal triglyceride levels in larvae expressing GFP or Dif in fat body, n = 18-20/group. **p = 0.0034 versus GFP. (**D**) Whole-animal triglyceride levels in larvae expressing GFP or Toll^10b^+Dif^RNAi^ in fat body, n = 8/group. Data are presented as mean ± SD. p values were determined by Student’s unpaired t test.

We investigated the mechanism for reduced triglyceride storage in the immune-activated fat body beginning with examining levels of circulating glucose, the substrate for *de novo* fatty acid and triglyceride synthesis. Trehalose, a disaccharide composed of two glucose molecules, is the major circulating sugar in fruit flies. Hemolymph trehalose and glucose levels were equivalent in larvae expressing GFP or Toll^10b^ in fat body (Fig 2A), suggesting that altered substrate availability does not account for reduced triglyceride storage. Another fate of glucose is storage as the branched polysaccharide glycogen. Surprisingly, whole-animal glycogen levels were increased in third instar larvae with active Toll signaling in fat body (Fig 2B). To better understand this phenotype, we measured glycogen in two tissues that are the major sites of glycogen storage in fly larvae, fat body and body wall muscles [26]. We observed a 3.7-fold increase in glycogen levels in fat body but no change in glycogen in the carcass, containing the body wall musculature, in animals with active fat body Toll signaling (Fig 2C). In white prepupae, glycogen levels were reduced from third instar levels and were equivalent in animals expressing GFP or Toll^10b^ in fat body (Fig 2D). These data show that triglyceride and glycogen storage are regulated differently by innate immune signaling and they suggest that a portion of the accumulated glycogen supports the final molt.

**Fig 2.**
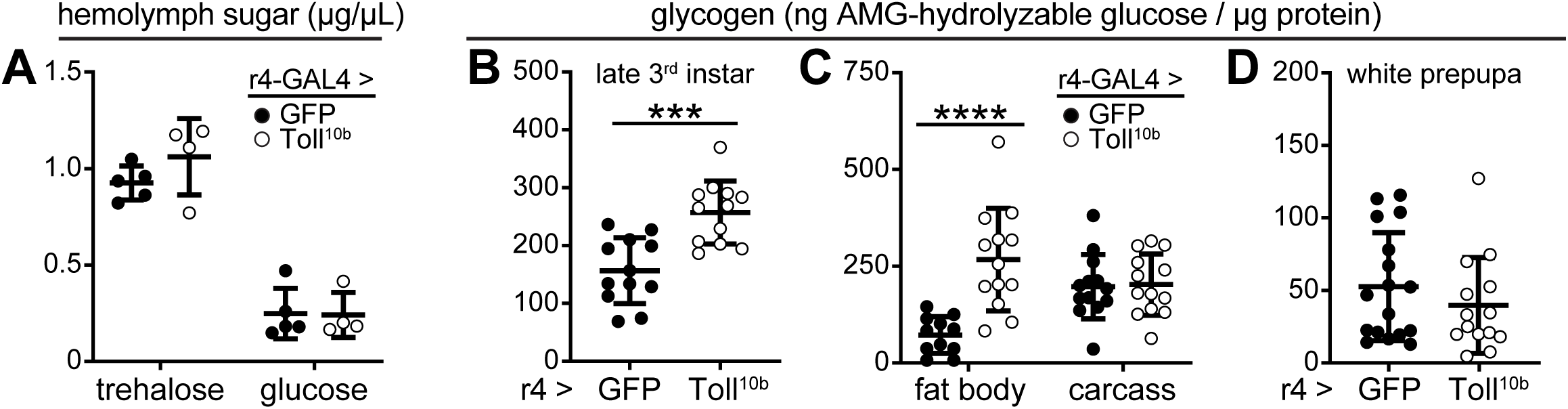
Toll signaling leads to increased fat body glycogen storage in the larval stage. (**A**) Hemolymph glucose and trehalose levels in mid-third instar larvae expressing GFP or Toll^10b^ in fat body under control of r4-GAL4, n = 4-5/group. (**B**) Late third instar whole-animal glycogen levels, normalized to protein, n = 12/group. ***p = 0.0002 versus GFP. (**C**) Glycogen levels in fat body and body wall, normalized to protein, from late third instar larvae expressing GFP or Toll^10b^ in fat body, n = 11-13/group, ****p < 0.0001 versus GFP. (**D**) White prepupal whole-animal glycogen levels, normalized to protein, n = 14-17/group. Data are presented as means ± SD. p values were determined by Student’s unpaired t test.

### Toll signaling negatively regulates dedicated steps of triglyceride synthesis

*De novo* lipogenesis is controlled in part by transcriptional regulation of genes encoding lipogenic enzymes [27]. Transcripts encoding *ATP citrate lyase* (*ATPCL*), *Acetyl-CoA carboxylase* (*ACC*) and *Fatty acid synthase 1* (*FASN1*), enzymes that synthesize fatty acids from glucose-derived citrate (Fig 3A), were expressed at equivalent levels in control and Toll^10b^-expressing fat bodies (Fig 3B). In contrast, Toll signaling in fat body led to 40-45% reductions in transcripts encoding *Lipin*, a phosphatidic acid phosphatase that synthesizes diacylglycerol from phosphatidic acid (Fig 3C), and *midway*, the *Drosophila* homolog of diacylglycerol acetyltransferase (DGAT), that carries out the final step in triglyceride synthesis (Fig 3D). Similarly, transgenic expression of Dif led to 24-33% reductions in *Lipin* and *midway* expression compared with GFP-expressing controls (Fig 3C and 3D).

**Fig 3.**
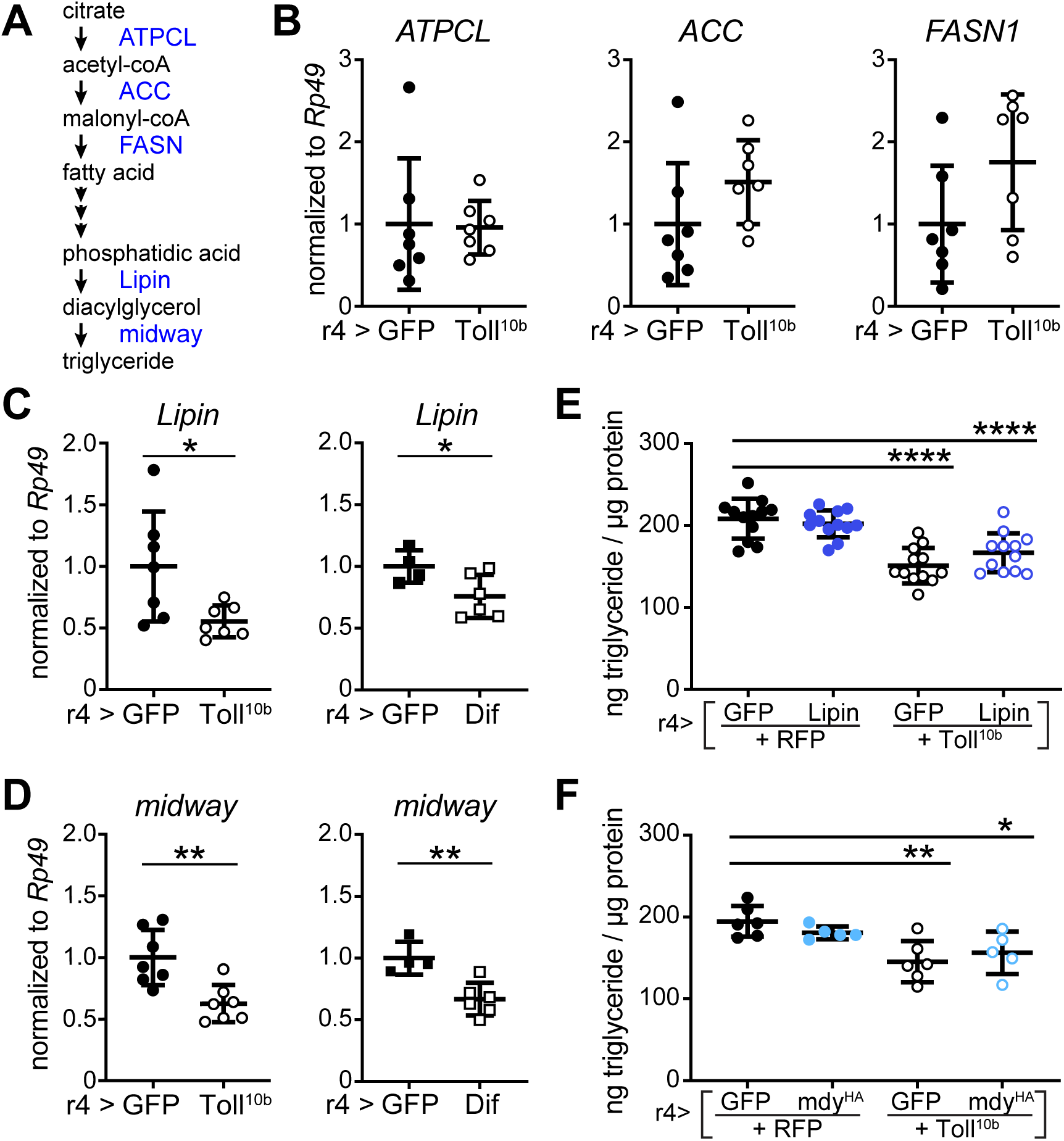
Reduced expression of *Lipin* and the DGAT homolog *midway* correlate with low triglyceride levels in larvae with active Toll signaling. (**A**) Schematic representation of the *de novo* lipogenic pathway leading to triglyceride synthesis. (**B-D**) Total RNA was extracted from late third instar larval fat bodies expressing the indicated transgenes under control of r4-GAL4. (**B**) *ATPCL, ACC*, and *FASN1* transcripts were measured by RT-qPCR and normalized to *Rp49*, n = 7/group. (**C**) *Lipin* mRNA levels in late third instar larval fat bodies expressing GFP or Toll^10b^ (n = 7/group. *p = 0.0259 versus GFP) or GFP or Dif (n = 4-6/group. *p = 0.0470 versus GFP). Transcripts were normalized to *Rp49*. (**D**) *midway* mRNA levels in late-third instar larval fat bodies expressing GFP or Toll^10b^ (n = 7/group. **p = 0.0031 versus GFP) or GFP or Dif (n = 4-6/group. **p = 0.0047 versus GFP). Transcripts were normalized to *Rp49*. (**E**) Triglyceride levels in late third instar larvae expressing Toll^10b^ with or without a wild type Lipin transgene in fat body, n = 12/group. ****p < 0.0001 versus RFP+GFP. (**F**) Triglyceride levels in late third instar larvae expressing Toll^10b^ with or without a wild type mdy^HA^ transgene in fat body, n = 5-6/group. *p = 0.0193, **p = 0.0020 versus RFP+GFP. Data are shown as means ± SD. p values were determined by Student’s t test (B-D) and one-way ANOVA with Dunnett’s multiple comparisons test (E,F).

Both Lipin and midway are necessary for triglyceride storage as knockdown of either in fat body or loss of midway in the whole animal decreased triglyceride storage (S2A-S2C Fig), consistent with previous findings [28-31]. Elevated fat body expression of Lipin or midway was not sufficient to increase triglyceride accumulation in otherwise wild type larvae (S2D-S2F Fig). However, fat body-specific expression of a UAS-midway transgene restored triglycerides in flies heterozygous for the *mdy*^*QX25*^ null mutation (S2G Fig).

We asked whether triglyceride storage could be rescued by forcing expression of either Lipin or midway in animals with active Toll signaling in fat body. Elevated Lipin failed to rescue low triglyceride levels in larvae co-expressing Toll^10b^ (Fig 3E). Similarly, co-expression of a wild type midway transgene with Toll^10b^ did not rescue triglyceride storage (Fig 3F). The failure of Lipin or midway expression to rescue low triglycerides in larvae with active innate immune signaling is consistent with the possibility that fatty acids are being diverted to another purpose in cells with active Toll signaling and that elevated flux through this second pathway could prevent triglyceride accumulation.

### Innate immune signaling induces phospholipid synthesis

Given that *ATPCL, ACC* and *FASN1* are expressed at normal levels in fat bodies with active Toll signaling, we considered that fatty acids produced by the actions of these enzymes might be used for purposes other than triglyceride storage during the immune response. A major fate of fatty acids in cells is phospholipid synthesis via the Kennedy pathway (Fig 4A). We assessed transcript levels of Kennedy pathway homologs in fat bodies expressing GFP or Toll^10b^ under control of r4-GAL4. We find elevated transcript levels of both the ethanolamine kinase *easily shocked* (*eas*) and the diacylglycerol ethanolaminephosphotransferase *CG7149* in fat bodies with active Toll signaling compared with controls (Fig 4B). *Phosphocholine cytidylyltransferase 1* (*Pcyt1*) was also induced by Toll signaling (Fig 4C), while the choline kinase homolog *CG2201* was inconsistently increased by fat body Toll pathway activation (compare data from independent experiments shown in Fig 4C and S3A Fig). Protein levels of eas and Pcyt1 were noticeably induced in fat bodies with active Toll signaling (Fig 4D and 4E). We observed that expression of the Toll pathway effector Dif phenocopied Toll^10b^, leading to increased transcript levels of *eas* and *Pcyt1* compared with fat bodies expressing GFP (Fig 4F and 4G). Other enzymes in the Kennedy pathway, such as *Phosphoethanolamine cytidylyltransferase* (*Pect*), the diacylglycerol cholinephosphotransferase *bb in a boxcar* (*bbc*) (refer back to Fig 4B and 4C), *Phosphocholine cytidylyltransferase 2* (*Pcyt2*) and the diacylglycerol ethanolaminephosphotransferase *CG33116* (S3B Fig) were not induced by Toll signaling. We note that our previously published RNA-Seq data [25] show much higher expression of *Pcyt1* compared with *Pcyt2* in larval fat body (S3C Fig).

**Fig 4.**
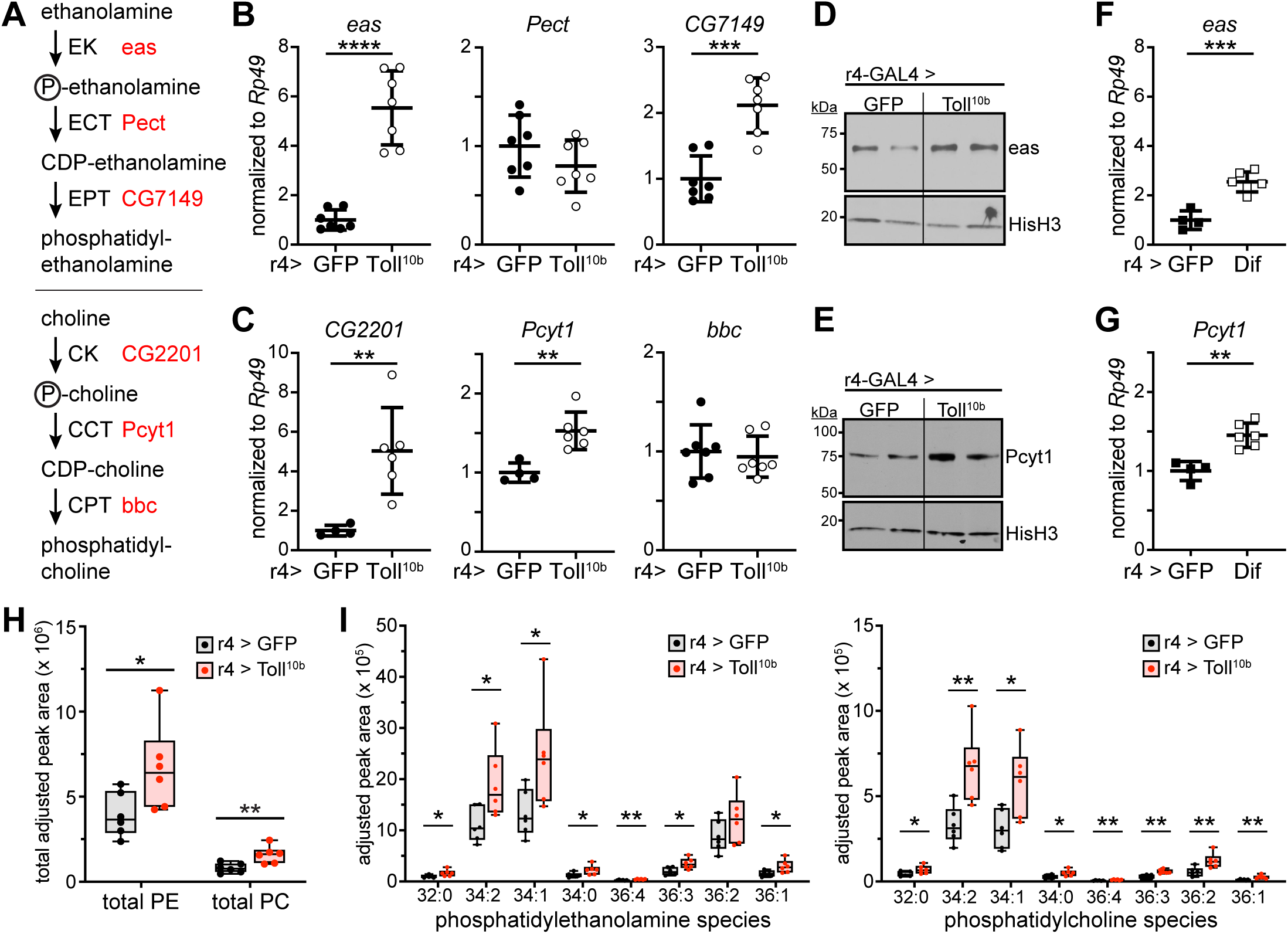
Toll signaling in fat body leads to a cascade of increased phospholipid synthesis enzymes and elevated membrane phospholipids. (**A**) Schematic representation of phosphatidylethanolamine (PE) and phosphatidylcholine (PC) synthesis via the Kennedy pathway. (**B**) Late third instar fat body levels of *eas, Pect*, and *CG7149* transcripts, normalized to *Rp49*, n = 7/group. ***p ≤ 0.0002, ****p = 0.0001 versus GFP. (**C**) Late third instar fat body levels of *CG2201, Pcyt1*, and *bbc* transcripts, normalized to *Rp49*. n = 7/group. **p ≤ 0.0071 versus GFP. (**D**,**E**) Western blot analysis of eas (D) and Pcyt1 (E) protein levels in fat bodies expressing GFP or Toll^10b^ under control of r4-GAL4. Histone H3 levels are shown as loading controls. (**F**,**G**) Late third instar fat body mRNA levels of *eas* (F) and *Pcyt1* (G) in fat bodies expressing GFP or Dif, n = 4-6/group. ***p = 0.0003 versus GFP (D), **p = 0.0011 versus GFP (E). Transcripts were normalized to *Rp49*. (**H**) Mass spectrometry analysis of total phosphatidylethanolamine levels (PE, left) and phosphatidylcholine levels (PC, right) in lipid extracts from late third instar larval fat bodies expressing GFP or Toll^10b^, n= 6/group. *p = 0.0417, **p = 0.0080 versus GFP. (**I**) Mass spectrometry analysis of PE (left) and PC species (right) in lipid extracts from late third instar larval fat bodies expressing GFP or Toll^10b^, n = 6/group. *p < 0.05, **p < 0.01 versus GFP. Data are shown as mean ± SD. p values were determined by Student’s unpaired t test.

To determine the functional consequences of increased *eas, CG7149, CG2201*, and *Pcyt1* expression in fat bodies with active Toll signaling, we assessed phospholipid levels in fat body. Using thin layer chromatography (TLC), we found increases in total phosphatidylethanolamine (PE) and phosphatidylcholine (PC) in fat bodies expressing Toll^10b^ compared with controls expressing GFP (S3D and S3E Fig). As a more sensitive assay, we used mass spectrometry to measure levels of the major PE and PC species in fat body [32-34]. In agreement with our TLC results, we found increased total levels of PE and PC in fat bodies expressing Toll^10b^ compared with controls (Fig 4H). We note that our TLC and mass spectrometry data show higher levels of PE than PC in both genotypes, as is true of membrane phospholipid composition in flies, and opposite to the pattern in mammals, where PC predominates [35]. Levels of most major PE and PC species were increased 1.5-to 2-fold in Toll^10b^-expressing fat bodies compared with controls (Fig 4I). These data suggest that increased expression of Kennedy pathway enzymes translates to substantial changes in phospholipid metabolism in response to immune signaling.

### Transcriptional control of Kennedy pathway enzymes in cells with active Toll signaling

We next investigated the role of two transcription factors – Sterol regulatory element binding protein (SREBP) and X box binding protein-1 (Xbp1) – in the elevated expression of phospholipid synthesizing enzymes in fat bodies with active Toll signaling. In flies, as in mammals, the transcription factor SREBP is a key regulator of *de novo* lipogenesis [36]. We find that fat bodies expressing Toll^10b^ exhibit elevated expression of SREBP mRNA and protein (S4A and S4B Fig). However, fat body-specific knockdown of SREBP did not affect either basal or Toll signaling-induced expression of *eas, CG7149*, or *CG2201*. Although basal expression of *Pcyt1* was reduced nearly three-fold by loss of SREBP, activation of Toll signaling led to 1.8-fold induction of *Pcyt1* in fat bodies regardless of SREBP levels (S4C Fig).

The unfolded protein response mediator Xbp1 regulates *de novo* lipogenesis of membrane phospholipids to permit ER expansion [37]. Xbp1 is activated by Ire1-mediated splicing [38,39], and fat bodies with active Toll signaling, induced by expression of Toll^10b^ or Dif, exhibit elevated levels of spliced *Xbp1* compared with controls (Fig 5A and 5B and S4D Fig). Loss of Dif in fat bodies expressing Toll^10b^ prevented the increase in spliced *Xbp1* (Fig 5B). *Xbp1* null mutants die as second instar larvae [40], and chronic knockdown of *Xbp1* in larval fat body via r4-GAL4 strongly inhibits whole-animal growth (data not shown). However, acute induction of both Toll^10b^ and Xbp1^RNAi^ transgenes in fat body using temperature-sensitive TubGAL80ts to restrict r4-GAL4 activity to a 24-hour period led to a near-complete loss of *Xbp1* transcripts (Fig 5A). Knocking down *Xbp1* blunted Toll^10b^-dependent induction of *eas* and blocked Toll^10b^-dependent induction of *CG7149* (Fig 5C). Acute loss of *Xbp1* in fat body reduced basal levels of *Pcyt1* transcripts, as was the case with chronic loss of SREBP (refer back to S4C Fig). Acute expression of Toll^10b^ induced *Pcyt1* 1.6-fold compared with controls, but only 1.2-fold in fat bodies with simultaneous loss of *Xbp1* (Fig 5D). We did not observe induction of *CG2201* in this experiment, perhaps due to the acute nature of Toll^10b^ expression (Fig 5D). Finally, acute loss of *Xbp1* blunted induction of *SREBP* and also the AMP *Drs* by Toll signaling; we did not observe changes in *Drs* when SREBP was manipulated (S4D and S4E Fig).

**Fig 5.**
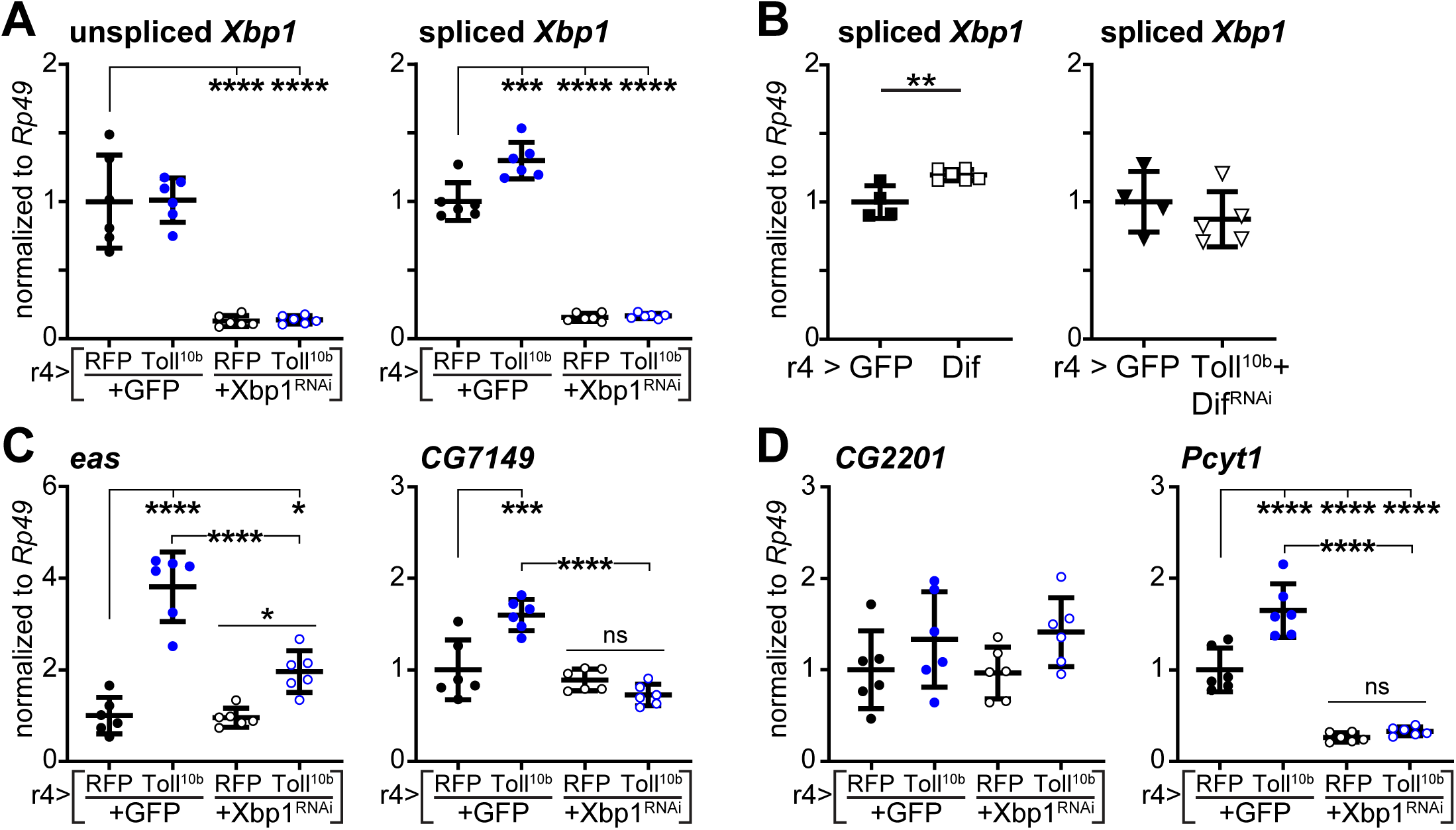
Toll signaling induces Kennedy pathway enzymes in a Xbp1-dependent manner. (**A**) Transcript levels of unspliced (left) and spliced (right) *Xbp1* were measured by RT-qPCR in late third instar larval fat bodies with GAL80ts-mediated induction of Toll^10b^ with or without Xbp1^RNAi^ for 24 hours at 30°C, n = 6/group. ***p = 0.0001 and ****p < 0.0001 versus fat bodies acutely expressing RFP+GFP. (**B**) Transcript levels of spliced *Xbp1* in fat bodies with r4-GAL4-driven expression of (left) GFP or Dif, n = 4-6/group, **p = 0.0049 versus GFP; or (right) GFP or Toll^10b^ + Dif^RNAi^, n = 4-5/group. (**C**, **D**) Transcript levels of indicated genes were measured by RT-qPCR in late third instar larval fat bodies with GAL80ts-mediated induction of Toll^10b^ with or without Xbp1^RNAi^ for 24 hours at 30°C, n = 6/group. **p = 0.0087, ***p = 0.0001 and ****p < 0.0001 versus fat bodies acutely expressing RFP+GFP. Data are presented as means ± SD. p values were determined by one-way ANOVA with Dunnett’s (A,C) or the Tukey-Kramer multiple comparisons test (D) and Student’s t test (B).

### Toll signaling leads to ER expansion

The induction of *Xbp1* splicing suggested that the ER unfolded protein response might be induced in fat bodies with active innate immune signaling. We therefore examined expression of canonical ER resident proteins that participate in the unfolded protein response and found significantly elevated expression of *binding immunoglobulin protein* (*BiP*), *protein disulfide isomerase* (*Pdi*), and *ER degradation enhancer mannosidase alpha-like 1* (*Edem1*) in response to chronic Toll signaling (Fig 6A).

**Fig 6.**
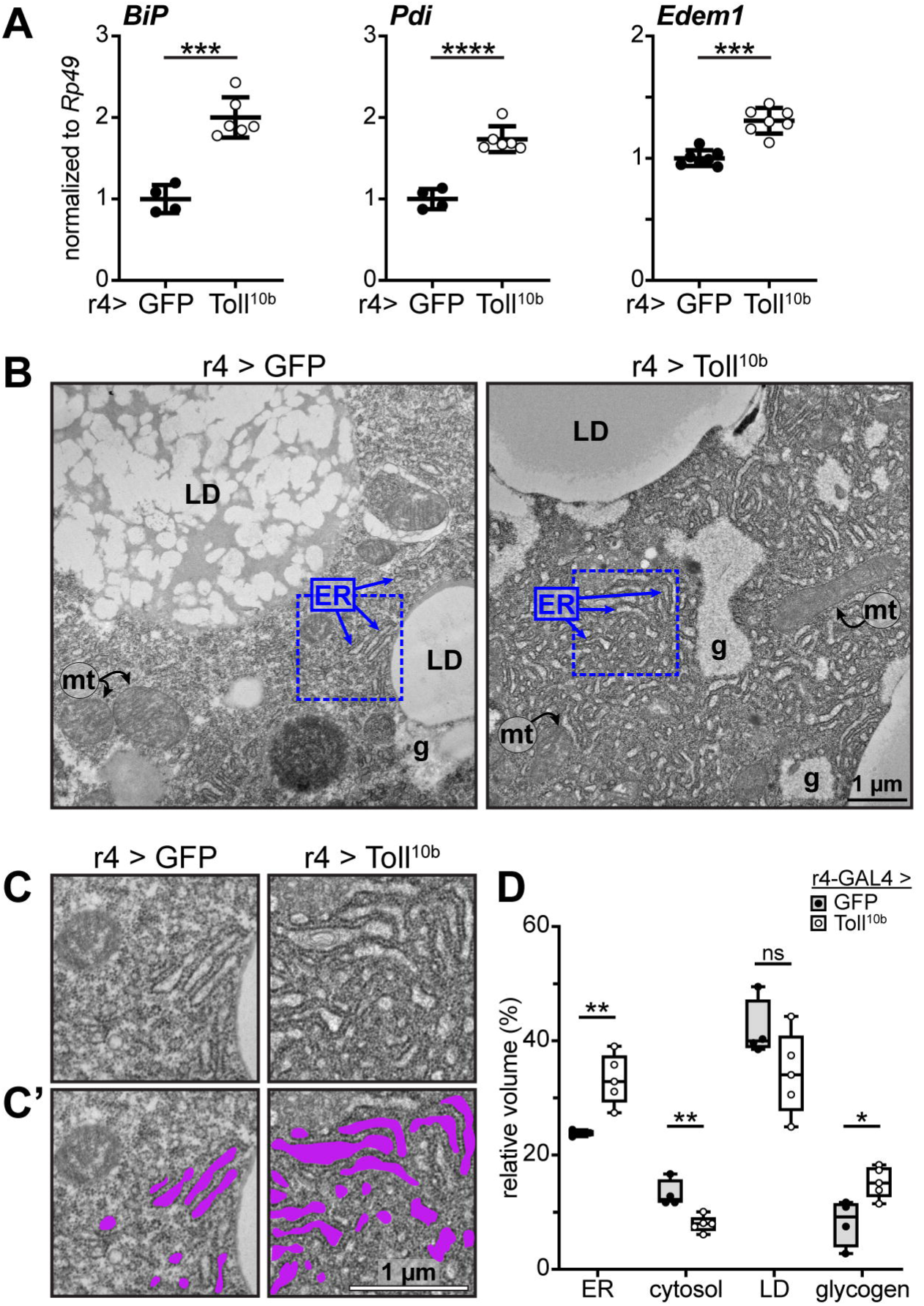
Chronic fat body Toll signaling leads to expansion of the endoplasmic reticulum. (**A**) Late third instar fat body levels of *BiP, Pdi* and *Edem1* were measured by RT-qPCR and normalized to *Rp49*, n = 4-7/group, ***p = 0.001, ****p < 0.0001 versus GFP. (**B**) Electron micrographs of fat body cells expressing GFP (left) or Toll^10b^ (right) under control of r4-GAL4. ER, endoplasmic reticulum; LD, lipid droplet; g, glycogen; mt, mitochondria. Dashed blue lines show areas enlarged in (C and C’). Scale bar, 1 µm. (**C**) Insets of electron micrographs outlined in blue dashed boxes in (B). (**C’**) Insets with ER structures highlighted in purple. Scale bar, 1µm. (**D**) Stereological analysis of organelle- and feature-specific volume density. Data points represent the average of volume densities measured from 5-15 images from a single fat body, n = 4-5 fat bodies/group. *p = 0.0175, **p ≤ 0.0044 versus GFP. Data are presented as means ± SD. p values were determined by Student’s t test (A,D).

We next used transmission electron microscopy to examine whether ER morphology is altered by innate immune signaling. We observed dilated ER in fat body cells with active Toll signaling compared with controls (Fig 6B-C’). To quantify this difference, we used stereological tools to compare organelles and features such as glycogen and cytosol between genotypes. This analysis revealed a 40% increase in the relative volume of ER with a simultaneous 40% decrease in the relative volume of organelle-free cytosol in response to Toll pathway activation. We find no significant difference in the relative volume of lipid droplets, but we observe an increase in glycogen volume in fat body cells expressing Toll^10b^ compared with controls (Fig 6D).

### AMP synthesis contributes to induction of Kennedy pathway enzymes

During the immune response to fungi or Gram-positive bacteria, the Toll signaling pathway drives synthesis and secretion of large quantities of AMPs via the classical ER-Golgi-secretory vesicle pathway [41]. Individual AMPs have been measured at levels close to 100 µM in hemolymph, which represents secretion of millions of individual proteins [16]. Acute activation of Toll signaling leads to 4-to 410-fold induction of 17 of the 37 AMPs encoded in the *Drosophila* genome (S5A Fig) [25]. We reasoned that ER expansion and phospholipid synthesis may serve the process of AMP production and secretion. To test this hypothesis, we asked whether induction of Kennedy pathway enzymes occurs downstream of AMP synthesis. The *Bom*^*Δ55C*^ mutation deletes a cluster of ten highly-induced genes, the bomanins, that are critical for survival during infection [42], and the *Drs*^*Δ7-17*^ mutation is a deletion of *Drs* [43]. Together, the genes encoded in the *Bom*^*55C*^ cluster and *Drs* account for 80% of the AMP transcripts that are induced by Toll signaling (Fig 7A). In a series of genetic manipulations to curtail AMP production, we drove Toll^10b^ in fat bodies of larvae carrying *Bom*^*Δ55C*^ and *Drs*^*Δ7-17*^ mutations in single or double homozygous combinations and measured expression of the Kennedy pathway enzymes *eas* and *Pcyt1*. The *Bom*^*Δ55C*^ and *Drs*^*Δ7-17*^ mutations, alone or in combination, led to the expected decreases in Toll^10b^ induced expression of the *Bom*^*55C*^ gene *BomS2* and *Drs* (Fig 7B). Unmanipulated AMPs such as *IM4* and *IM14* were still induced by Toll^10b^ in *Bom*^*Δ55C*^ and *Drs*^*Δ7-17*^ single or double mutants (S5B Fig). In larval fat bodies expressing GFP, mutation of *Bom*^*Δ55C*^ and *Drs*^*Δ7-17*^ did not alter low, basal levels of *eas* or *Pcyt1*. However, *Bom*^*Δ55C*^; *Drs*^*Δ7-17*^ double homozygotes expressing Toll^10b^ exhibited only a 1.6-fold induction of *eas* and a 1.3-fold induction of *Pcyt1*. In contrast, in larvae with a full complement of AMPs, Toll signaling elicited a 4.1-fold induction of *eas* and a 2.1-fold induction of *Pcyt1* (Fig 7C and 7D). These data suggest that changes in Kennedy pathway enzymes occur downstream of AMP synthesis.

**Fig 7.**
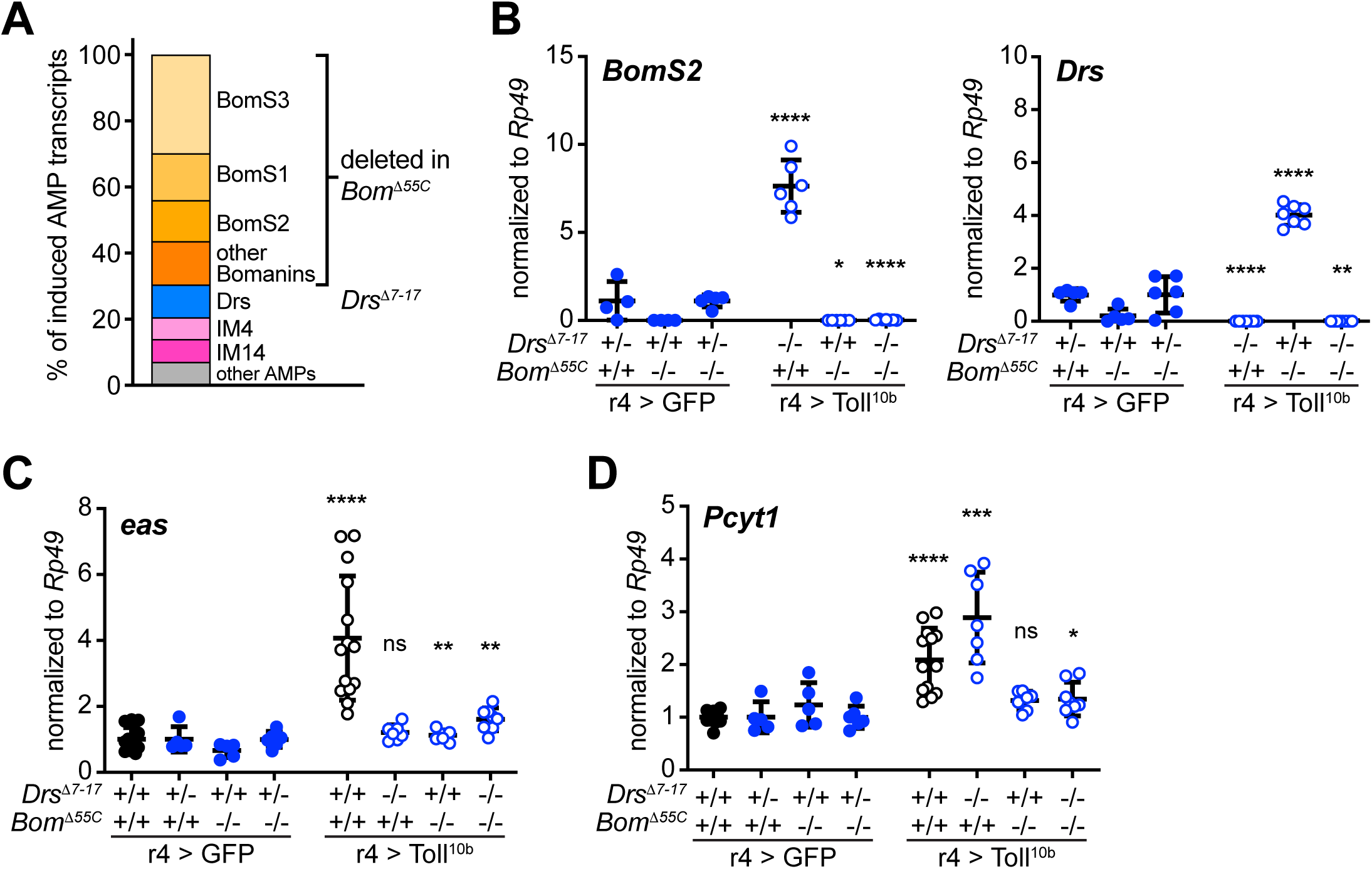
Induction of antimicrobial peptide synthesis contributes to regulation of eas and Pcyt1 by Toll signaling. (**A**) Toll signaling induces expression of 17 AMPs. AMPs clustered on chromosome 2 (deleted in the *Bom*^*Δ55C*^ mutant) represent 70% of the AMP transcripts induced by Toll^10b^ expression. Drs, deleted in the *Drs*^*Δ7-17*^ mutant, represents 10% of the induced AMP transcripts. (**B**) Transcript levels of *BomS2* (left) and *Drs* (right), normalized to *Rp49*, in fat bodies of late third instar larvae expressing GFP (closed symbols) or Toll^10b^ (open symbols) under r4-GAL4 control. Animals were wild type, heterozygous, or homozygous for *Drs*^*Δ7-17*^ and *Bom*^*Δ55C*^ as indicated, n = 5-8/group. *p = 0.0291, **p = 0.0015, ****p < 0.0001 versus GFP-expressing controls with the same *Drs*^*Δ7-17*^ and *Bom*^*Δ55C*^ genotypes. Note that GFP-expressing controls are heterozygous for *Drs*^*Δ7-17*^ while Toll^10b^-expressing larvae are homozygous for *Drs*^*Δ7-17*^. (**C**, **D**) Transcript levels of *eas* (C) and *Pcyt1* (D), normalized to *Rp49*, in fat bodies of late third instar larvae expressing GFP (closed symbols) or Toll^10b^ (open symbols) under r4-GAL4 control. Animals were wild type, heterozygous, or homozygous for *Drs*^*Δ7-17*^ and *Bom*^*Δ55C*^ as indicated, n = 5-8/group (blue open and closed symbols) and n = 11-14/group for animals expressing GFP or Toll^10b^ in fat body on a wild type background (black open and closed symbols). *p ≤ 0.0409, **p ≤ 0.032, ***p = 0.0009, ****p < 0.0001. Data are presented as means ± SD. p values were determined by Student’s t test.

## Discussion

Here we show that chronic Toll pathway activity acts in a tissue-autonomous manner to reduce triglyceride storage and increase phospholipid levels in the *Drosophila* larval fat body, an organ analogous to mammalian liver and adipose tissue that also serves innate immune functions. This decrease in triglycerides is not due to a decrease in availability of the substrate glucose; indeed, larvae with active Toll signaling exhibit a transient increase in fat body glycogen storage compared with controls. Instead, transcript levels of *Lipin* and the DGAT homolog *midway*, enzymes that regulate the final steps of triglyceride synthesis, are selectively decreased in fat bodies expressing active Toll receptors, mirroring decreased triglyceride levels. A second major fate of fatty acids is phospholipid synthesis, and we discovered that fat bodies with active Toll signaling exhibit tissue-autonomous increases in levels of major PE and PC species. Kennedy pathway enzymes that synthesize PE and PC are increased at the transcript and protein levels in larvae with active Toll signaling in fat body, in parallel with increased phospholipid levels. This increase in phospholipid enzyme transcripts is dependent on the transcription factor Xbp1, an important regulator of ER biogenesis and the unfolded protein response. Accordingly, morphological analysis shows expanded and dilated ER in fat bodies with active Toll signaling. In response to activation of the Toll pathway, the fat body synthesizes and secretes massive quantities of antimicrobial peptides to defend against pathogenic Gram-positive bacteria and fungi. We show that deletion of AMP genes blunts the Toll-dependent induction of key enzymes that synthesize phospholipids, suggesting that elevated synthesis of these secreted proteins, and possibly ER stress, contributes to induction of the Kennedy pathway. These data suggest that the animal induces expression of enzymes that synthesize membrane phospholipids in order to sustain production of AMPs at a high level within the secretory pathway.

Our results demonstrate a reciprocal relationship between triglyceride and phospholipid levels in the *Drosophila* larval fat body. Activating innate immune signaling in this organ leads to a decrease in triglyceride storage and increases in PE and PC species. Such a tradeoff between triglyceride storage and membrane phospholipid levels is observed in other contexts. In S2 cells, in adult flies, in *C. elegans*, and in mammals, loss of Kennedy pathway enzymes leads to elevated triglyceride levels [44-48]. In contrast, manipulations that reduce lipin and DGAT activity directly or indirectly by disrupting Torsin function lead to elevated phospholipid levels and reduced triglyceride synthesis [29,49,50]. These studies and ours suggest that cells tightly regulate the balance between stored neutral lipids and membrane phospholipids.

Reduced triglyceride storage in animals with active fat body Toll signaling may be due, at least in part, to changes in expression of two key lipogenic enzymes, the phosphatidic acid phosphatase Lipin and the DGAT homolog midway. However, we were unable to rescue impaired triglyceride storage caused by Toll signaling by restoring Lipin or midway expression in fat body. A possible explanation for this result is that Toll signaling may regulate Lipin or midway not only at the level of gene expression but also at the level of enzyme activity. Alternatively, Toll activation may disrupt subcellular localization of Lipin such that transgenic expression of either Lipin or midway would be insufficient to promote triglyceride accumulation. Indeed, flies with loss of the ER membrane protein Torsin exhibit an increase in nuclear Lipin and elevated triglyceride levels [29], and in mammals, insulin signaling promotes lipin activity by changing its subcellular distribution [51]. Our previous work shows that Toll signaling in fat body inhibits insulin signaling, and that restoring insulin signaling rescues triglyceride storage [24]. Therefore, Toll signaling might be expected to dominantly inhibit Lipin function by altering its subcellular distribution. Finally, the failure of forced Lipin or midway expression to restore triglyceride accumulation in fat bodies with active Toll signaling may be due to an increase in flux of lipid intermediates into the phospholipid synthesis pathway.

Major species of the membrane phospholipids PE and PC accumulate in response to Toll pathway activation. Our results suggest that increased phospholipid levels are likely due to increased synthesis of PE and PC because Kennedy pathway enzymes are increased at transcript and protein levels in fat bodies expressing Toll^10b^. We find elevated protein levels of eas and Pcyt1, enzymes that carry out rate-limiting first and second steps, respectively, in the PE and PC synthesis pathways [52,53]. Loss-of-function mutations in *eas* and *Pcyt1* lead to reduced levels of PE and PC [54,55], and these mutations lead to defects in neuronal morphogenesis and excitability [56,57], oocyte development [58], and cardiac function [47]. Our data suggest that these enzymes also contribute to the immune response in *Drosophila*.

Our investigation into the transcriptional mechanism underlying increased expression of genes encoding Kennedy pathway enzymes led us to discover that Xbp1 participates in this response. Xbp1 is a transcription factor that is activated by splicing of its transcript when high secretory demand leads to an abundance of misfolded proteins in the ER. Xbp1 serves as an essential mediator of the unfolded protein response that relieves ER stress, and its role in the whole animal extends to supporting function of cells with high secretory capacity. For example, during differentiation of naïve B cells into immunoglobulin-secreting plasma cells, Xbp1 is induced and is required for maximal immunoglobulin secretion [11]. Additionally. *Xbp1* is required for the membrane phospholipid synthesis and ER expansion that accompany plasma cell differentiation [9,10,59]. In mouse fibroblasts, forced expression of spliced *Xbp1* is sufficient to induce PC and PE synthesis via the Kennedy pathway and to drive ER biogenesis [37,60]. Our data suggest that Xbp1-dependent induction of phospholipid synthesis and ER expansion occur in response to Toll signaling in *Drosophila*. We show that *Xbp1* is spliced in response to Toll signaling, and that Xbp1 is necessary for maximal induction of *eas, CG7149* and *Pcyt1* in response to acute activation of fat body Toll signaling. Toll receptor activation leads to a massive induction of AMP secretion, and we find that relieving secretory demand by deletion of AMP genes blunts induction of *eas* and *Pcyt1*. Our results suggest that immune effector production leading to ER stress-induced synthesis of phospholipids and subsequent ER biogenesis that supports immune function is an ancient and conserved component of the immune response.

Our data are compatible with either a linear or a parallel relationship between AMP production and Kennedy pathway enzyme expression. It is possible that the tremendous secretory demand induced by Toll signaling leads to ER stress, subsequent activation of Xbp1, downstream expression of phospholipid synthesis enzymes that generate PE and PC to expand the ER and reduce stress. Indeed, deletion of AMP genes blunts induction of *eas* and *Pcyt1* by Toll signaling. However, it is also possible that the Toll pathway induces Kennedy pathway enzymes, in a Xbp1-dependent manner, in parallel with elevated AMP expression, thereby anticipating upcoming secretory demand. Our data would support such a model if a feedback mechanism would allow levels of phospholipid-synthesizing enzymes to be titrated to the actual secretory demand. Distinguishing between these models will require careful analysis of the early time course of AMP and Kennedy pathway enzyme transcription in response to Toll pathway activation.

The shift from nutrient storage to membrane phospholipid synthesis induced by Toll signaling likely has both immediate and long-term consequences for animal survival. The lipid metabolic switch accompanies ER expansion, and biogenesis of ER is predicted to sustain secretion of AMPs during infection. On the other hand, it is critical to tightly regulate ER function during infection. Unrestrained Ire1 activity and Xbp1 splicing in *sigma-1 receptor* knockout mice leads to elevated proinflammatory cytokine production and increased rates of sepsis in response to LPS treatment [61]. An increase in host PE and PC synthesis would lead to increased consumption of ethanolamine and choline. These molecules can be used as carbon and nitrogen sources for bacteria [62,63], therefore increased host Kennedy pathway activity may also lead to reduced growth for some species of bacteria. A clear disadvantage of shifting fatty acids from nutrient storage to phospholipid synthesis in response to Toll signaling in flies is the large decrease in stored energy available for metamorphosis and early adult life. Animals with active Toll signaling enter the pupal stage with 50% less triglyceride compared with control animals. Chronic activation of Toll signaling leads to a 80% decrease in adult viability, with most animals dying at the pharate adult stage [24]. This decrease in viability is likely multifactorial, but impaired triglyceride levels likely lead to reduced metabolic energy available to complete metamorphosis as well as a decrease in the pool of triglyceride that is needed to waterproof the adult cuticle [64].

Taken together, our results show that Toll signaling leads to a profound change in lipid metabolism that supports immune function but also hinders the ability of the animal to store energy. Our work raises the question of how animals balance the metabolic demands of infection with the capacity to survive periods of reduced nutrient availability. Furthermore, it will be of interest to determine whether chronic changes in lipid metabolism induced by innate immune signaling underlie pathologies of inflammatory diseases.

## Materials and Methods

### *Drosophila* stocks and husbandry

Flies were raised on food containing 7.8% molasses, 2.4% yeast, 4.6% cornmeal, 0.3% propionic acid, and 0.1% methylparaben (Archon Scientific, Durham, NC). Except where noted, experiments were performed using mid-to late-third instar larvae (96-108 h after egg lay). The following stocks were obtained from the Bloomington *Drosophila* Stock Center (Bloomington, IN): UAS-RFP (#30556), UAS-GFP (2nd (#1521) and 3rd (#1522) chromosome insertions), UAS-Dif^RNAi^ (#30513), *mdy*^*QX25*^ (#5095), *mdy*^*EY07280*^ (#20167), UAS-midway^RNAi^ (#65963), UAS-Lipin^RNAi^ (#63614), UAS-Xbp1^RNAi^ (#36755), and r4-GAL4 (#33832). UAS-SREBP^RNAi^ (#37640) were obtained from the Vienna *Drosophila* Resource Center. Other flies used were: UAS-Toll^10b^ [65], UAS-Dif [66], UAS-Lipin [30], *Bom*^*Δ55C*^ [42], and *Drs*^*Δ7-17*^ [43]. Full genotypes of flies are listed in S1 Table.

### Construction of UAS-HA.mdy transgenic flies

The full-length *midway* (*mdy*) cDNA was amplified by PCR from clone LD33582 (*Drosophila* Genomics Resource Center, Bloomington, IN) using gene-specific primers engineered to contain an amino-terminal HA tag (see S2 Table). PCR products were cloned into pENTR (Invitrogen) and validated by sequencing. Gateway cloning (Invitrogen) was used to generate pUAST-HA.mdy. This construct was injected into *Drosophila* embryos at Rainbow Transgenics (Camarillo, CA). Standard genetics was used to map and generate balanced transgenic lines.

### Triglyceride and protein measurements

Whole larvae or dissected organs were flash frozen on dry ice. Samples were sonicated three times for 10 seconds each time in 140mM NaCl, 50mM Tris-HCl, pH 7.4, 0.1% Triton X-100 with protease inhibitors (Roche). Following clearing by centrifugation at 4°C, supernatants were transferred to new tubes. Triglyceride (Liquicolor Test, Stanbio) and protein (BCA assay, Pierce) were measured in each sample, and triglyceride levels were normalized to protein levels.

### Hemolymph trehalose and glucose

Hemolymph was collected on ice from mid-third instar larvae (hemolymph from 8-10 larvae pooled per sample). Endogenous trehalase was destroyed by heating hemolymph diluted in PBS at 70°C for 20 min. The sample was split in half, 1 mU trehalase (Sigma-Aldrich, T8778) was added to one tube, and both were incubated at 37°C for 2 h. Glucose was measured in both samples (GAGO20 kit, Sigma-Aldrich). Trehalose was calculated by subtracting glucose values in trehalase-free samples from glucose values in trehalase-treated samples, and then dividing by two as trehalose is a dimer of glucose. Trehalose and glucose values were normalized to hemolymph volume.

### Glycogen and free glucose measurements

Whole larvae or dissected organs were flash frozen on dry ice. Samples were homogenized using a Kontes pestle in 0.1M NaOH. Following clearing by centrifugation at 4°C, samples were incubated for 1h at 37°C with 0.2M NaOAc, pH 4.8 with or without 5mg/mL amyloglucosidase (Sigma-Aldrich, A7420). Samples were then incubated for 10 min at room temperature with assay buffer including glucose oxidase (0.25 U/mL, Sigma-Aldrich, G7141) and horseradish peroxidase (0.17 U/mL, Sigma-Aldrich, P8250) and 20µM Amplex Red (Invitrogen, A36006). Fluorescence was measured (excitation/emission maxima = 535/587 nm) using an Infinite 200 PRO plate reader (Tecan). Glycogen was calculated as the amount of amyloglucosidase-hydrolyzed glucose, and free glucose is the amount of glucose measured in the sample without hydrolysis via amyloglucosidase. Glycogen and free glucose levels were normalized first to glucose standards (Fisher CAS 50-99-7) and then to protein levels (BCA assay, Pierce).

### Western blot analysis and antibodies

Fat bodies (n = 4-6 pooled/sample) were sonicated in lysis buffer (2% SDS, 60 mM Tris-HCl, pH 6.8) with phosphatase and protease inhibitors (Roche). Protein concentration was measured using a BCA assay (Pierce). Equal amounts of protein (10-40 µg/lane) were separated by SDS-PAGE, transferred to nitrocellulose, blocked in 3% milk in 1X TBS with 0.2% Tween 20 (TBS-T) and blotted overnight at 4°C with primary antibodies diluted in 1% milk in TBS-T. Following multiple washes in TBS-T, secondary antibodies were incubated in 1% milk in TBS-T for 2h at room temperature, washed again, incubated with ECL (Pierce) and exposed to film. Antibodies used were: rabbit anti-human Histone H3, and rabbit anti-HA (4499 and 3724, Cell Signaling Technology); rabbit anti-*Drosophila* easily shocked [67], guinea pig anti-*Drosophila* Pcyt1 [29], mouse anti-human SREBP1 (SC-13551, Santa Cruz Biotechnology), rabbit anti-GFP (A11122, Invitrogen), goat anti-rabbit HRP and goat anti-mouse HRP (111-035-003 and 115-035-003, Jackson ImmunoResearch), and goat anti-guinea pig HRP (6090-05, SouthernBiotech).

### Quantitative RT-PCR

Total RNA was extracted from late third instar fat bodies (n = 4-6 pooled/sample) using a Direct-zol RNA MicroPrep kit (Zymo Research). DNAse-treated total RNA (1 µg) was used to generate cDNA using a High-Capacity cDNA Reverse Transcription kit (Thermo Fisher Scientific). Gene expression was measured using gene-specific primers. Quantitative PCR reactions were performed on 10-20 ng cDNA using SYBR Select Master Mix (Thermo Fisher Scientific) with a Bio-Rad CFX Connect Real-Time PCR Detection System. Relative amounts of transcripts were calculated using the comparative Ct method with *Rp49* as a reference gene [68]. Gene-specific primer sequences are listed in S2 Table.

### Thin Layer Chromatography

Late third instar larval fat bodies (n = 8 pooled/ sample) were sonicated in 1X TBS. Fat body lysates underwent lipid extraction using a modified Bligh-Dyer method (chloroform: methanol: 0.2 M NaCl, ratio 2:2:1) [69]. Organic extracts were dried and resuspended in 30µL of chloroform: methanol (1:1) for quantitation by thin layer chromatography. Using a Hamilton syringe (#701), 15-20µL of resuspended extracts were spotted on the bottoms of silica gel plates (Millipore HPTLC Silica Gel Glass plates 105631) at starting points drawn in pencil. Standards were phosphatidylcholine (3µL at 1µg/µL; egg L-α-phosphatidylcholine Avanti Polar Lipids, 131601) and phosphatidylethanolamine (3µL at 1µg/µL; egg L-α-phosphatidylethanolamine Avanti Polar Lipids, 840021). During sample loading, plates were placed beneath a pure argon gas flow to allow for rapid drying and minimal oxidation of organic extracts. Silica gel plates were placed in a glass TLC tank with elution buffer specifically to elute phospholipids (chloroform: methanol: acetic acid: water in a ratio of 75: 35: 6: 2) and covered with glass lid. Plates were left in elution buffer until liquid reached 1cm from the top of the plate. Plates were removed, air dried, and left in a separate glass TLC tank in a fume hood with a small beaker of iodine crystals overnight. For analysis of silica gel plates, plates with iodine stain were scanned and images underwent densitometry analysis in ImageJ to measure the area of each PE and PC spot, recognized based on the location and migration pattern of the corresponding standard. Areas of lipid spots were normalized to total protein in lysates (measured by BCA assay, Pierce) and volumes of lysates used in the initial organic extraction.

### Liquid chromatography and mass spectrometry

Larval fat bodies (n = 6 pooled/sample) were sonicated in ultrapure water and protein was measured (BCA assay, Pierce). Fat body lysates (normalized to 40 µg protein) underwent lipid extraction using a modified Bligh-Dyer method (chloroform: methanol: ultrapure water, ratio 2:2:1), adding 5µg 1,2-dinonadecanoyl-sn-glycero-3-phosphocholine (DNPC, Avanti Polar Lipids, 850320) to control for extraction efficiency. Organic extracts were dried and resuspended in chloroform: methanol (1:1) for quantification by liquid chromatography-coupled mass spectrometry. Lipids were separated on a EVO C18 column (Kinetex 5µm, 100 x 4.6 mm, Phenomenex), using a binary gradient consisting of Solvent A (69% methanol, 31% water with 10mM ammonium acetate) and Solvent B (50% methanol, 50% isopropanol with 10mM ammonium acetate) as the mobile phases. Phosphatidylcholine (PC) and phosphatidylethanolamine (PE) species identification and relative quantitation was achieved by liquid chromatography-linked electrospray ionization mass spectrometry on a 4000 Q Trap triple-quadrupole mass spectrometer (AB Sciex). Identification of phospholipids of interest, corresponding to PC and PE species with acyl chain lengths previously described to be abundant in larval fat body [32-34], was performed in positive ion mode via multiple reaction monitoring [70].

### Electron Microscopy

Larval fat bodies were dissected and fixed initially in 2% glutaraldehyde, 2.5% formaldehyde, 0.1M Na cacodylate, pH 7.4 (30 min at room temperature followed by 1h on ice). They were then post-fixed in 1% OsO4, 0.1M Na cacodylate, pH 7.4 (15 min on ice and 45 min at room temperature), washed three times for 5 min in 0.1M Na acetate, pH 6.0, and then stained in-block overnight at room temperature in 0.5% uranyl acetate in the same buffer. The samples were then dehydrated in acetone (70, 95, and 100%), exchanged from acetone into Epon resin, and embedded in fresh Epon. 70 nm sections from embedded fat bodies were mounted on copper grids, stained sequentially with uranyl acetate and lead citrate, and examined by transmission electron microscopy.

### Stereology

All images for stereology were taken in an unbiased manner at 6000X. Three different blocks of each genotype (r4>GFP or r4>Toll^10b^) that each contained fat bodies from three to four larvae were used for quantitative analysis. Biological replicates refer to sets of adjacent sections cut from separate blocks or from different depths of the same block to expose different fat body tissue. STEPanizer software (Java) was used for stereology analysis [71]. In order to measure relative volumes of features within sections, images were analyzed using nine-line tile pairs per image [72]. Point counts were made to determine relative volume of organelles comprising the entire cytoplasmic landscape in each image. Batch mode was used to analyze images from each block for each genotype.

### Quantitation and statistical analysis

Statistical parameters including exact sample sizes (with n referring to biological replicates), data plotted (typically mean ± SD), exact p values, and statistical tests used are reported in Figure Legends. Statistical analyses were performed using Graphpad Prism 8. Data were analyzed by Student’s t test or by one-way ANOVA with the Tukey-Kramer or Dunnett’s multiple comparisons test.

## Acknowledgements

We thank Young Jun Lee (University of Virginia) and Stacey Criswell (Advanced Microscopy Facility, University of Virginia) for help with experiments. We thank Tony Ip (University of Massachusetts, Worcester), Michael Lehmann (University of Arkansas), Steven Wasserman (UCSD) and Takayuki Kuraishi (Kanazawa University, Japan) for flies, Thomas Preat (École des Neurosciences, Paris) for the eas antibody, Rose Goodchild (KU Leuven, Belgium) for the Pcyt1 antibody. We thank Thurl Harris, Janet Cross (both University of Virginia) and members of the Bland lab, especially Miyuki Suzawa, for discussions. We thank the Bloomington *Drosophila* Stock Center and *Drosophila* Genomics Resource Center (both Bloomington, IN) and the Vienna *Drosophila* Resource Center (Vienna, Austria) for reagents. This work was supported by NIH Grant R01DK099601 to M.L.B. and Ruth L. Kirschstein NRSA Predoctoral Fellowship F31DK118879 to B.A.M., NIH Grant P01 HL120840 to N.L., NIH MSTP T32 GM007267 to S.Y. and NIH Pharmacological Sciences Training Grant T32 GM007055-44 to B.A.M. and S.Y.

## Author Contributions

### Conceptualization

Brittany Martínez, Michelle Bland

### Formal Analysis

Brittany Martínez, Scott Yeudall, Rosalie Hoyle, David Castle, Michelle Bland

### Funding Acquisition

Norbert Leitinger, Michelle Bland

### Investigation

Brittany Martínez, Scott Yeudall, Rosalie Hoyle, David Castle, Michelle Bland

### Methodology

Brittany Martínez, Scott Yeudall, David Castle, Michelle Bland

### Validation

Brittany Martínez, Rosalie Hoyle, Michelle Bland

### Visualization

Brittany Martínez, Michelle Bland

### Writing – original draft

Brittany Martínez, Michelle Bland

### Writing – review & editing

Brittany Martínez, Scott Yeudall, Rosalie Hoyle, David Castle, Norbert Leitinger, Michelle Bland

## Supporting information

**S1 Fig.**
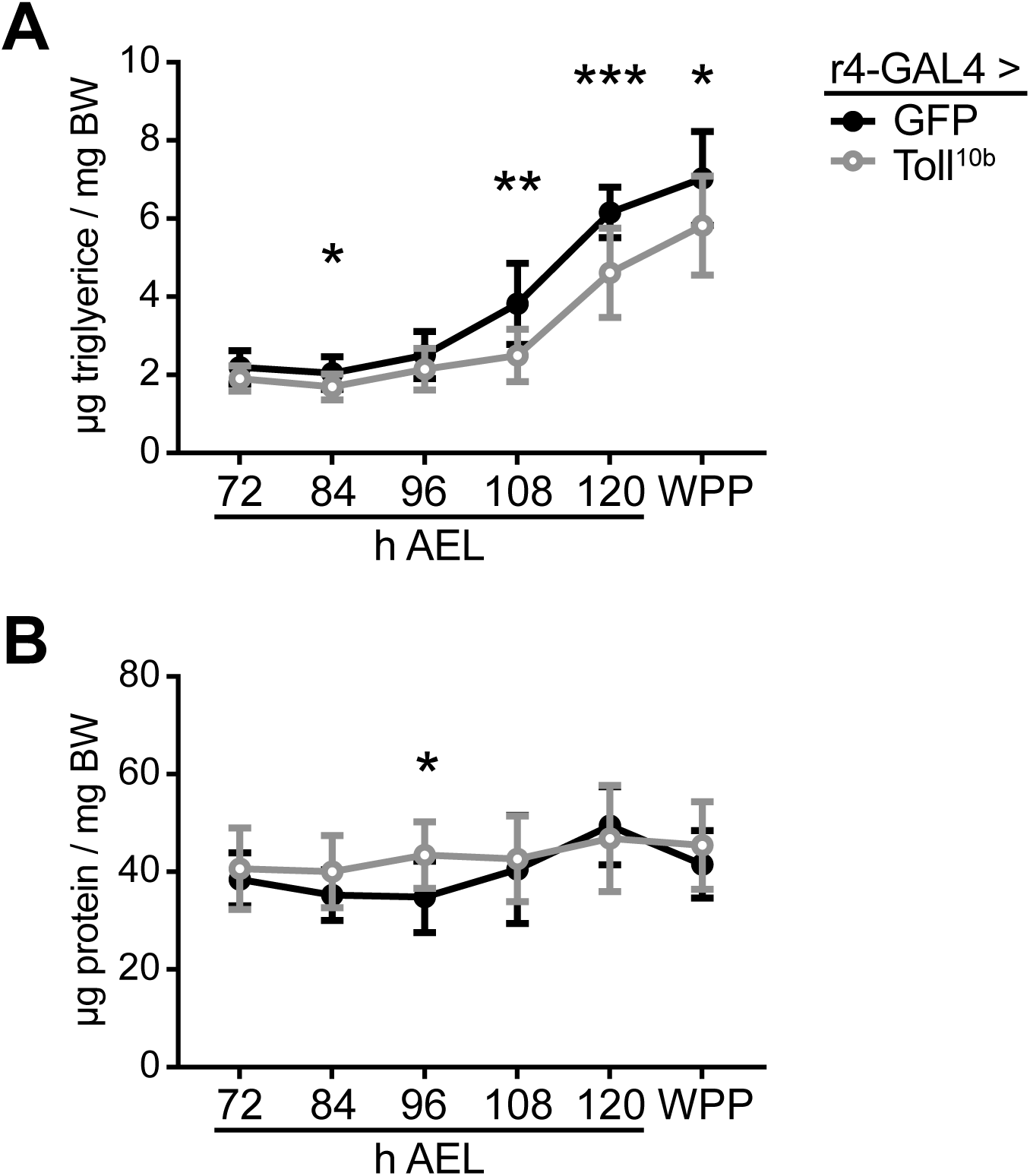
Toll signaling in the larval fat body reduces triglyceride storage throughout the third instar. (**A**) Whole-animal triglyceride levels throughout the third instar (72-120 hours after egg lay (h AEL)) and in white prepupae (WPP) were normalized to body weight, n = 10-11/group. *p ≤ 0.0373, **p = 0.0033, and ***p = 0.0009 versus GFP. (**B**) Whole-animal protein levels throughout the third instar (72-120 h AEL) and in white prepupae (WPP) were normalized to body weight, n = 10-11/group. *p = 0.0117 versus GFP. Data are presented as mean ± SD. p values were determined by Student’s unpaired t test.

**S2 Fig.**
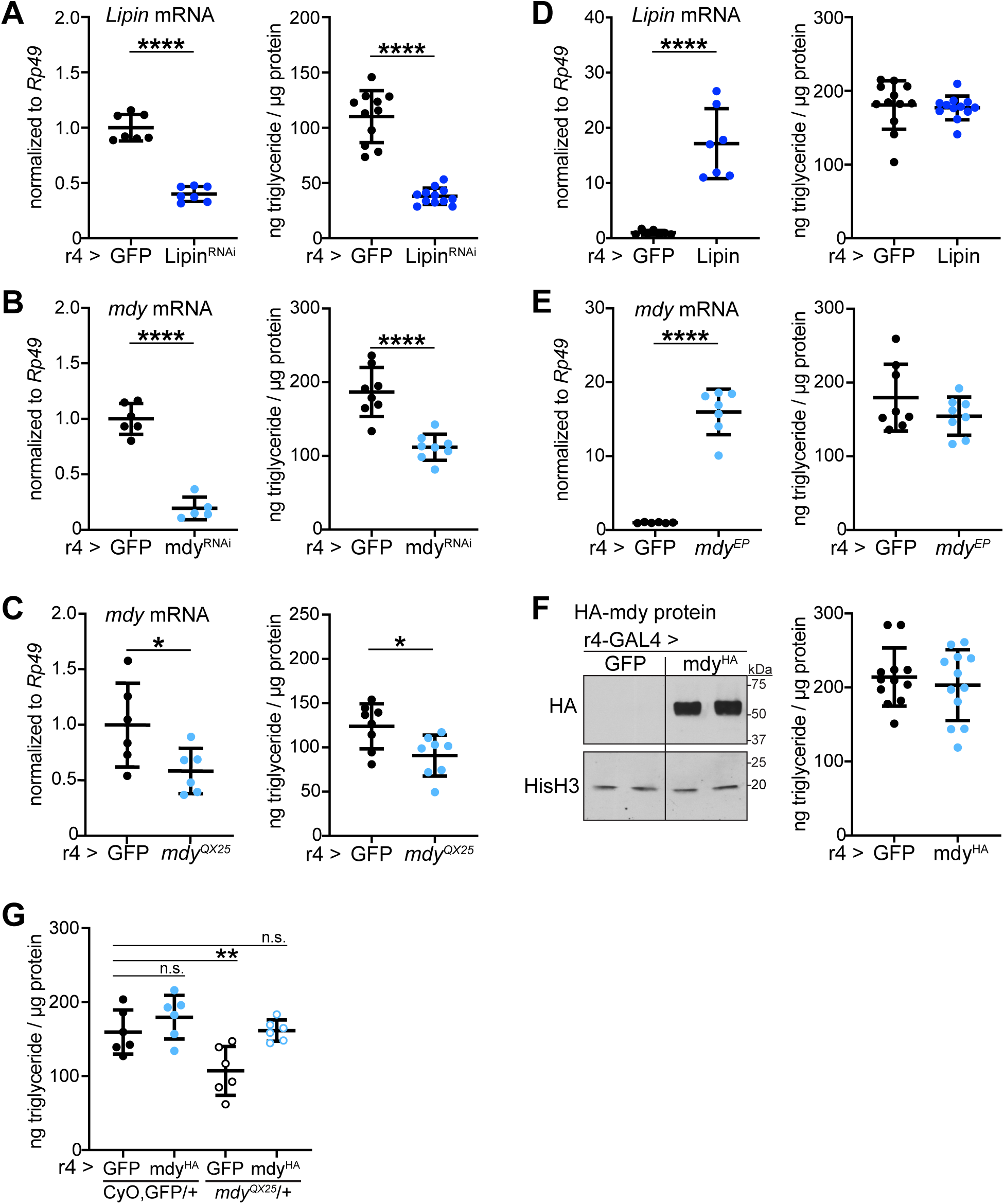
Lipin and midway are necessary for fat body triglyceride storage. For A-E, left panels: late third instar fat body levels of *Lipin* or *midway* mRNA, normalized to *Rp49*; right panels: late third instar whole-animal triglyceride levels, normalized to protein, in larvae of the indicated genotypes. (**A**) *Lipin* mRNA (n = 7/group) and triglycerides (n = 11-12/group) in larvae expressing GFP or Lipin^RNAi^ in fat body using r4-GAL4. ****p < 0.0001 versus GFP. (**B**) *midway* mRNA (n = 5-6/group) and triglycerides (n = 8/group) in larvae expressing GFP or mdy^RNAi^ in fat body. ****p < 0.0001 versus GFP. (**C**) *midway* mRNA (n = 6/group) and triglycerides (n = 8/group) in UAS-GFP/+; r4-GAL4/+ and *mdy*^*QX25*^/+; r4-GAL4/+ larvae. *p ≤ 0.0412 versus UAS-GFP/+; r4-GAL4/+. (**D**) *Lipin* mRNA (n = 7/group) and triglycerides (n = 12/group) in larvae expressing GFP or wild type Lipin in fat body. ****p < 0.0001 versus GFP. (**E**) *midway* mRNA (n = 6-7/group) and triglycerides (n = 8/group) in larvae with r4-GAL4 driven expression of UAS-GFP or *mdy*^*EY07280*^ in fat body. ****p < 0.0001 versus GFP. (**F**) Left: Western blot of HA-tagged midway transgene expression in fat bodies expressing GFP or wild type, HA-tagged midway (UAS-mdy^HA^) under r4-GAL4 control (HA, top). Histone H3 (bottom) is shown as a loading control. Right: whole-animal triglycerides in larvae expressing GFP or mdy^HA^ in fat body, n = 12/group. (**G**) Triglyceride levels in CyO, GFP/+ and *mdy*^*QX25*^/+ larvae expressing GFP or mdy^HA^ in fat body, n = 6/group. **p = 0.0097 versus CyO, GFP/+; r4-GAL4/UAS-GFP. Data are presented as means ± SD. p values were determined by Student’s t test (A-F) and one-way ANOVA with Dunnett’s multiple comparison test (G).

**S3 Fig.**
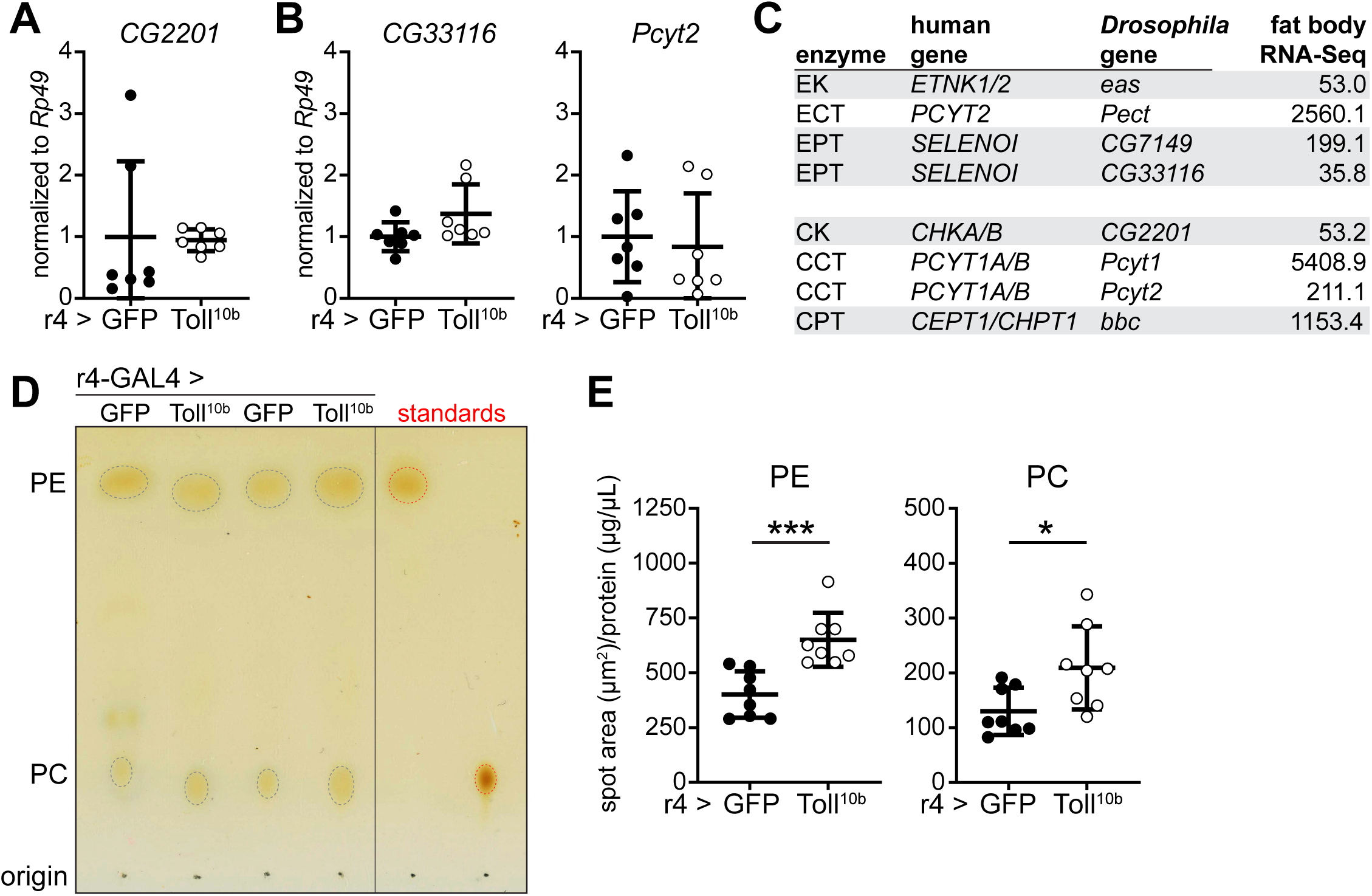
Expression of Kennedy pathway enzymes and elevated levels of membrane phospholipids in fat bodies with active Toll signaling. (**A**) Late third instar fat body levels of *CG2201* transcripts, normalized to *Rp49*, n = 7/group. (**B**) Late third instar fat body levels of *CG33116* and *Pcyt2* transcripts, normalized to *Rp49*, n = 7/group. (**C**) Basal levels of Kennedy pathway enzymes in control fat bodies, measured by RNA-Sequencing. Normalized read count data shown are from Suzawa et al., 2019. (**D**) Representative image of TLC separation of PE (top) and PC (bottom) from pooled larval fat bodies (n = 8/sample) expressing GFP or Toll^10b^ driven by r4-GAL4. Standards are outlined in red. (**E**) Area of PE and PC from densitometry of iodine-stained TLC plates, normalized to total protein in lysates, n = 8 measurements/genotype from three separate experiments. ***p = 0.0007 versus GFP (PE) and *p = 0.0146 versus GFP (PC). Data are presented as means ± SD. p values were determined by Student’s t test.

**S4 Fig.**
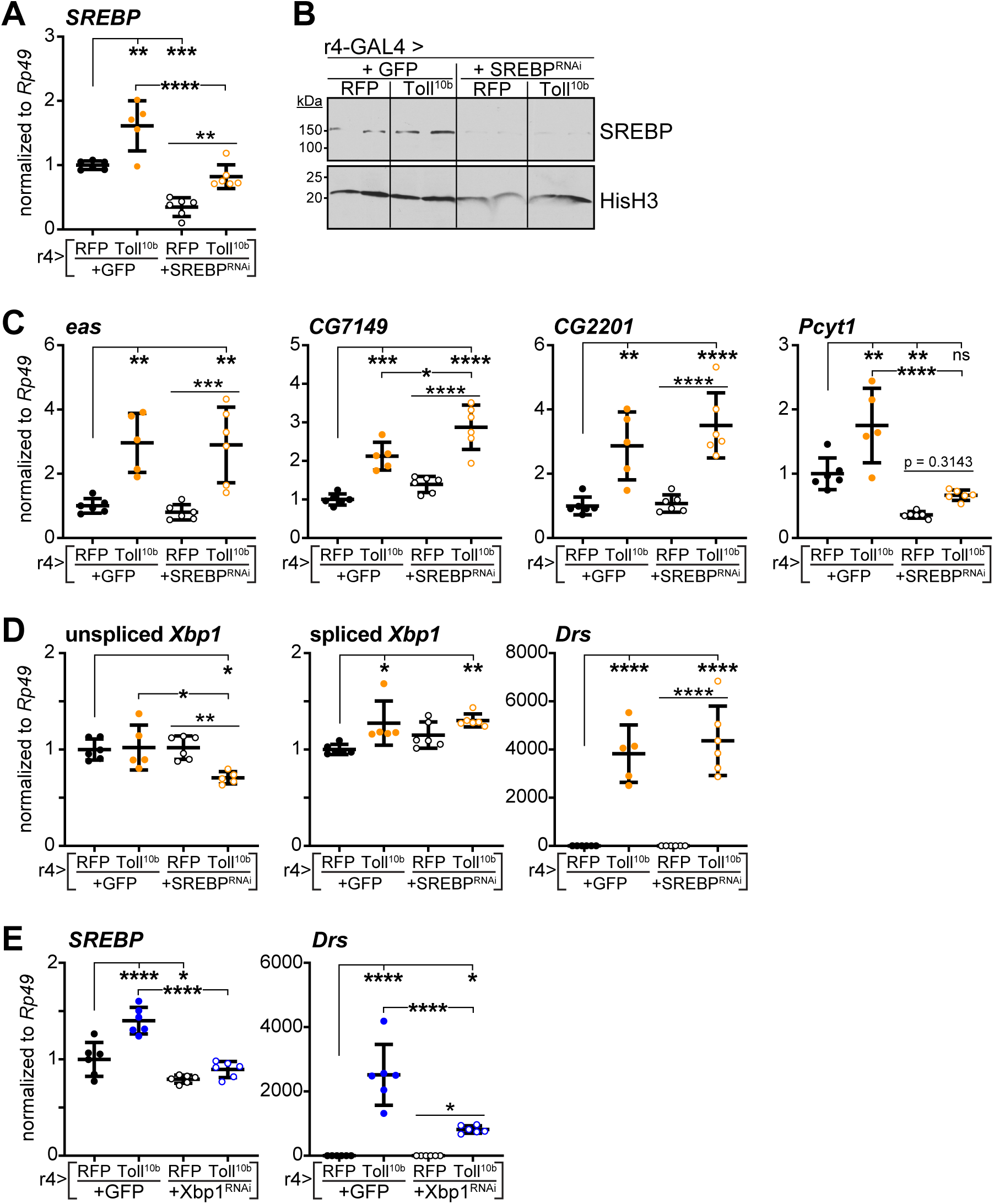
Toll signaling induces Kennedy pathway enzymes independently of SREBP. (**A**) Late third instar fat body *SREBP* transcript levels, normalized to *Rp49*, in fat bodies expressing RFP or Toll^10b^ with GFP or SREBP^RNAi^ under control of r4-GAL4, n = 5-6/group. **p ≤ 0.0069, ***p = 0.0003, ****p < 0.0001 versus RFP+GFP. (**B**) Western blot of SREBP (top) and Histone H3 (bottom) in fat bodies of the indicated genotypes. (**C**) Late third instar fat body mRNA levels of Kennedy pathway enzymes *eas, CG7149, CG2201* and *Pcyt1*, normalized to *Rp49*, in fat bodies of the indicated genotypes, n = 5-6/group. *p = 0.0139, **p ≤ 0.0073, ***p ≤ 0.0006, and ****p < 0.0001 versus RFP+GFP. (**D**) Late third instar fat body mRNA levels of unspliced and spliced *Xbp1* and *Drs*, normalized to *Rp49*, in fat bodies of the indicated genotypes, n = 5-6/group. *p ≤ 0.0155, **p ≤ 0.0096, and ****p < 0.0001 versus RFP+GFP. (**E**) Transcript levels of *SREBP* and *Drs* were measured by RT-qPCR in late third instar larval fat bodies with GAL80ts-mediated induction of Toll^10b^ with or without Xbp1^RNAi^ for 24 hours at 30°C, n = 6/group. *p ≤ 0.0366 and ****p < 0.0001 versus fat bodies acutely expressing RFP+GFP. Data are presented as means ± SD. p values were determined by one-way ANOVA with the Tukey-Kramer multiple comparisons test (A, C-E).

**S5 Fig.**
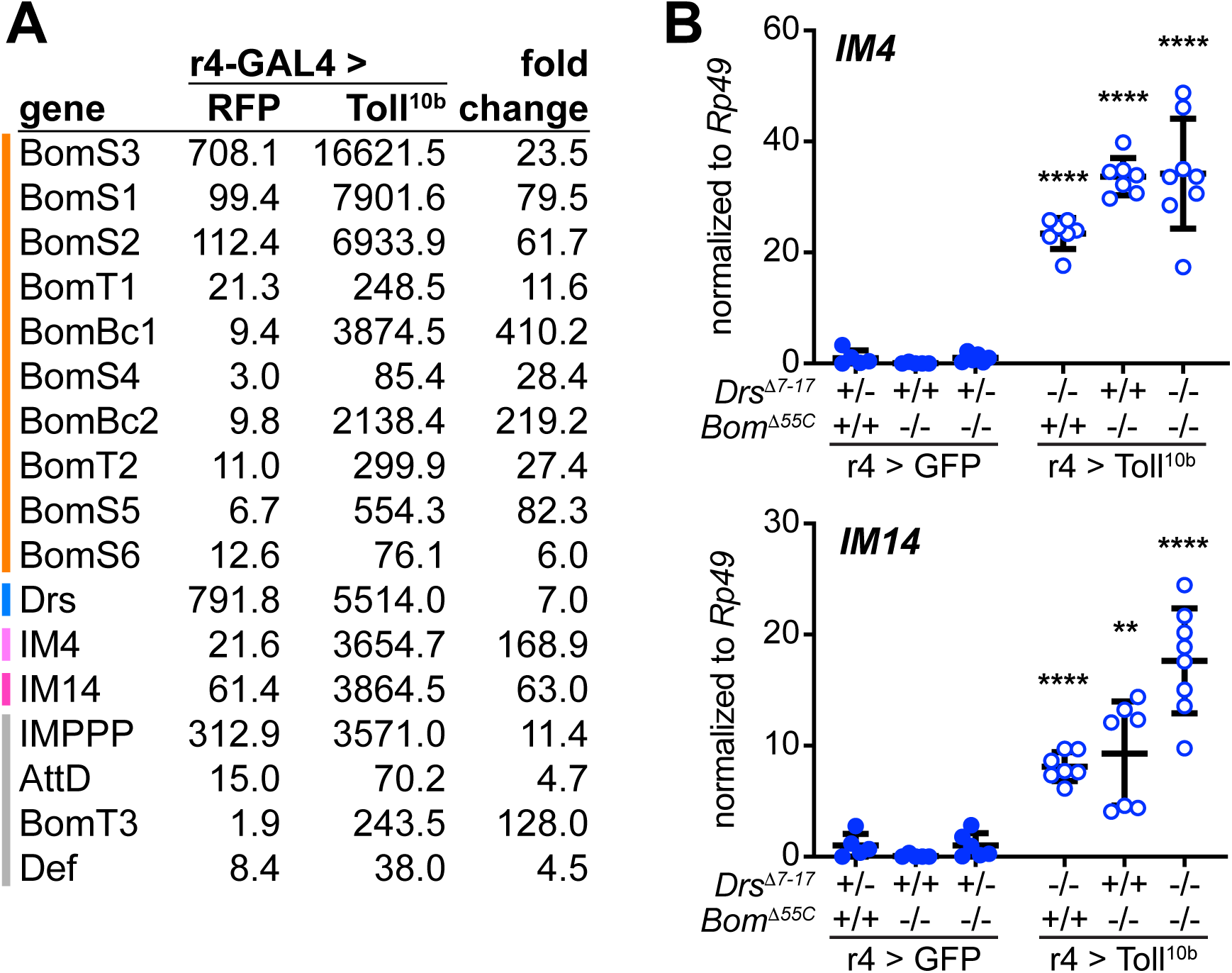
Untargeted immune molecules are induced in fat bodies with *Bom*^*Δ55C*^ and *Drs*^*Δ7-17*^ mutations. (**A**) Expression levels of AMPs in control fat bodies expressing RFP and in immune-activated fat bodies expressing Toll^10b^ under r4-GAL4 control. Normalized read count data from RNA-sequencing published in Suzawa et al., 2019 are shown. AMPs clustered on chromosome 2 (deleted in the *Bom*^*Δ55C*^ mutant, orange line) represent 70% of the AMP transcripts induced by Toll^10b^ expression. Drs, deleted in the *Drs*^*Δ7-17*^ mutant, represents 10% of the induced AMP transcripts. (**B**) Transcript levels of *IM4* (top) and *IM14* (bottom), normalized to *Rp49*, in fat bodies of late third instar larvae expressing GFP (closed symbols) or Toll^10b^ (open symbols) under r4-GAL4 control. Animals were wild type, heterozygous, or homozygous for *Drs*^*Δ7-17*^ and *Bom*^*Δ55C*^ as indicated, n = 5-8/group. **p = 0.0015, ****p < 0.0001 versus GFP-expressing controls with the same *Drs*^*Δ7-17*^ and *Bom*^*Δ55C*^ genotypes. Note that GFP-expressing controls are heterozygous for *Drs*^*Δ7-17*^ while Toll^10b^-expressing larvae are homozygous for *Drs*^*Δ7-17*^. Data are presented as means ± SD. p values were determined by Student’s t test.

**S1 Table.** Genotypes of *Drosophila melanogaster* used in this study.

**S2 Table.** Sequences of oligonucleotides used in this study.

## Notes

### Competing Interest Statement

The authors have declared no competing interest.

## References

1. Corrigan JJ, Fonseca MT, Flatow EA, Lewis K, Steiner AA. Hypometabolism and hypothermia in the rat model of endotoxic shock: independence of circulatory hypoxia. J Physiol (Lond). 2014;592: 3901–3916. doi:10.1113/jphysiol.2014.277277

2. Ganeshan K, Nikkanen J, Man K, Leong YA, Sogawa Y, Maschek JA, et al. Energetic Trade-Offs and Hypometabolic States Promote Disease Tolerance. Cell. 2019;177: 399–413.e12. doi:10.1016/j.cell.2019.01.050

3. Chang C-H, Curtis JD, Maggi LB, Faubert B, Villarino AV, O’Sullivan D, et al. Posttranscriptional control of T cell effector function by aerobic glycolysis. Cell. 2013;153: 1239–1251. doi:10.1016/j.cell.2013.05.016

4. Gleeson LE, Sheedy FJ, Palsson-McDermott EM, Triglia D, O’Leary SM, O’Sullivan MP, et al. Cutting Edge: Mycobacterium tuberculosis Induces Aerobic Glycolysis in Human Alveolar Macrophages That Is Required for Control of Intracellular Bacillary Replication. J Immunol. American Association of Immunologists; 2016;196: 2444–2449. doi:10.4049/jimmunol.1501612

5. Krawczyk CM, Holowka T, Sun J, Blagih J, Amiel E, DeBerardinis RJ, et al. Toll-like receptor-induced changes in glycolytic metabolism regulate dendritic cell activation. Blood. 2010;115: 4742–4749. doi:10.1182/blood-2009-10-249540

6. Krejčová G, Danielová A, Nedbalová P, Kazek M, Strych L, Chawla G, et al. Drosophila macrophages switch to aerobic glycolysis to mount effective antibacterial defense. Elife. 2019;8: 102. doi:10.7554/eLife.50414

7. Everts B, Amiel E, Huang SC-C, Smith AM, Chang C-H, Lam WY, et al. TLR-driven early glycolytic reprogramming via the kinases TBK1-IKKε supports the anabolic demands of dendritic cell activation. Nat Immunol. Nature Publishing Group; 2014;15: 323–332. doi:10.1038/ni.2833

8. van Anken E, Romijn EP, Maggioni C, Mezghrani A, Sitia R, Braakman I, et al. Sequential waves of functionally related proteins are expressed when B cells prepare for antibody secretion. Immunity. 2003;18: 243–253.

9. Fagone P, Sriburi R, Ward-Chapman C, Frank M, Wang J, Gunter C, et al. Phospholipid biosynthesis program underlying membrane expansion during B-lymphocyte differentiation. J Biol Chem. American Society for Biochemistry and Molecular Biology; 2007;282: 7591–7605. doi:10.1074/jbc.M608175200

10. McGehee AM, Dougan SK, Klemm EJ, Shui G, Park B, Kim Y-M, et al. XBP-1-deficient plasmablasts show normal protein folding but altered glycosylation and lipid synthesis. The Journal of Immunology. 2009;183: 3690–3699. doi:10.4049/jimmunol.0900953

11. Reimold AM, Iwakoshi NN, Manis J, Vallabhajosyula P, Szomolanyi-Tsuda E, Gravallese EM, et al. Plasma cell differentiation requires the transcription factor XBP-1. Nature. 2001;412: 300–307. doi:10.1038/35085509

12. Martinon F, Chen X, Lee A-H, Glimcher LH. TLR activation of the transcription factor XBP1 regulates innate immune responses in macrophages. Nat Immunol. 2010;11: 411–418. doi:10.1038/ni.1857

13. Tian Y, Pate C, Andreolotti A, Wang L, Tuomanen E, Boyd K, et al. Cytokine secretion requires phosphatidylcholine synthesis. J Cell Biol. Rockefeller University Press; 2008;181: 945–957. doi:10.1083/jcb.200706152

14. Buchon N, Silverman N, Cherry S. Immunity in Drosophila melanogaster--from microbial recognition to whole-organism physiology. Nat Rev Immunol. 2014;14: 796–810. doi:10.1038/nri3763

15. Valanne S, Wang J-H, Rämet M. The Drosophila Toll signaling pathway. J Immunol. 2011;186: 649–656. doi:10.4049/jimmunol.1002302

16. Fehlbaum P, Bulet P, Michaut L, Lagueux M, Broekaert WF, Hetru C, et al. Insect immunity. Septic injury of Drosophila induces the synthesis of a potent antifungal peptide with sequence homology to plant antifungal peptides. J Biol Chem. 1994;269: 33159–33163.

17. Hanson MA, Dostálová A, Ceroni C, Poidevin M, Kondo S, Lemaitre B. Synergy and remarkable specificity of antimicrobial peptides in vivo using a systematic knockout approach. Elife. 2019;8: 511. doi:10.7554/eLife.44341

18. Arrese EL, Soulages JL. Insect Fat Body: Energy, Metabolism, and Regulation. Annu Rev Entomol. 2010;55: 207–225. doi:10.1146/annurev-ento-112408-085356

19. Lehmann M. Endocrine and physiological regulation of neutral fat storage in Drosophila. Mol Cell Endocrinol. 2018;461: 165–177. doi:10.1016/j.mce.2017.09.008

20. Dionne MS, Pham LN, Shirasu-Hiza M, Schneider DS. Akt and FOXO dysregulation contribute to infection-induced wasting in Drosophila. Curr Biol. 2006;16: 1977–1985. doi:10.1016/j.cub.2006.08.052

21. Péan CB, Schiebler M, Tan SWS, Sharrock JA, Kierdorf K, Brown KP, et al. Regulation of phagocyte triglyceride by a STAT-ATG2 pathway controls mycobacterial infection. Nat Commun. Nature Publishing Group; 2017;8: 14642. doi:10.1038/ncomms14642

22. Franchet A, Niehus S, Caravello G, Ferrandon D. Phosphatidic acid as a limiting host metabolite for the proliferation of the microsporidium Tubulinosema ratisbonensis in Drosophila flies. Nat Microbiol. Nature Publishing Group; 2019;4: 645–655. doi:10.1038/s41564-018-0344-y

23. DiAngelo JR, Bland ML, Bambina S, Cherry S, Birnbaum MJ. The immune response attenuates growth and nutrient storage in Drosophila by reducing insulin signaling. Proc Natl Acad Sci USA. 2009;106: 20853–20858. doi:10.1073/pnas.0906749106

24. Roth SW, Bitterman MD, Birnbaum MJ, Bland ML. Innate Immune Signaling in Drosophila Blocks Insulin Signaling by Uncoupling PI(3,4,5)P3 Production and Akt Activation. CellReports. ElsevierCompany; 2018;22: 2550–2556. doi:10.1016/j.celrep.2018.02.033

25. Suzawa M, Muhammad NM, Joseph BS, Bland ML. The Toll Signaling Pathway Targets the Insulin-like Peptide Dilp6 to Inhibit Growth in Drosophila. CellReports. 2019;28: 1439–1446.e5. doi:10.1016/j.celrep.2019.07.015

26. Yamada T, Habara O, Kubo H, Nishimura T. Fat body glycogen serves as a metabolic safeguard for the maintenance of sugar levels in Drosophila. Development. Oxford University Press for The Company of Biologists Limited; 2018;145: dev158865. doi:10.1242/dev.158865

27. Wang Y, Viscarra J, Kim S-J, Sul HS. Transcriptional regulation of hepatic lipogenesis. Nat Rev Mol Cell Biol. Nature Publishing Group; 2015;16: 678–689. doi:10.1038/nrm4074

28. Beller M, Bulankina AV, Hsiao H-H, Urlaub H, Jäckle H, Kühnlein RP. PERILIPIN-Dependent Control of Lipid Droplet Structure and Fat Storage in Drosophila. Cell Metabolism. Elsevier Inc; 2010;12: 521–532. doi:10.1016/j.cmet.2010.10.001

29. Grillet M, Dominguez Gonzalez B, Sicart A, Pöttler M, Cascalho A, Billion K, et al. Torsins Are Essential Regulators of Cellular Lipid Metabolism. Dev Cell. 2016;38: 235–247. doi:10.1016/j.devcel.2016.06.017

30. Schmitt S, Ugrankar R, Greene SE, Prajapati M, Lehmann M. Drosophila Lipin interacts with insulin and TOR signaling pathways in the control of growth and lipid metabolism. J Cell Sci. 2015;128: 4395–4406. doi:10.1242/jcs.173740

31. Ugrankar R, Liu Y, Provaznik J, Schmitt S, Lehmann M. Lipin is a central regulator of adipose tissue development and function in Drosophila melanogaster. Mol Cell Biol. 2011;31: 1646–1656. doi:10.1128/MCB.01335-10

32. Carvalho M, Sampaio JL, Palm W, Brankatschk M, Eaton S, Shevchenko A. Effects of diet and development on the Drosophila lipidome. Mol Syst Biol. 2012;8. doi:10.1038/msb.2012.29

33. Guan XL, Cestra G, Shui G, Kuhrs A, Schittenhelm RB, Hafen E, et al. Biochemical Membrane Lipidomics during Drosophila Development. Dev Cell. Elsevier Inc; 2013;24: 98–111. doi:10.1016/j.devcel.2012.11.012

34. Palm W, Sampaio JL, Brankatschk M, Carvalho M, Mahmoud A, Shevchenko A, et al. Lipoproteins in Drosophila melanogaster—Assembly, Function, and Influence on Tissue Lipid Composition. P Kühnlein R, editor. PLoS Genet. 2012;8: e1002828. doi:10.1371/journal.pgen.1002828.g008

35. Dobrosotskaya IY, Seegmiller AC, Brown MS, Goldstein JL, Rawson RB. Regulation of SREBP processing and membrane lipid production by phospholipids in Drosophila. Science (New York, NY). American Association for the Advancement of Science; 2002;296: 879–883. doi:10.1126/science.1071124

36. Horton JD, Goldstein JL, Brown MS. SREBPs: activators of the complete program of cholesterol and fatty acid synthesis in the liver. J Clin Invest. American Society for Clinical Investigation; 2002;109: 1125–1131. doi:10.1172/JCI15593

37. Sriburi R, Jackowski S, Mori K, Brewer JW. XBP1: a link between the unfolded protein response, lipid biosynthesis, and biogenesis of the endoplasmic reticulum. J Cell Biol. 2004;167: 35–41. doi:10.1083/jcb.200406136

38. Cox JS, Walter P. A novel mechanism for regulating activity of a transcription factor that controls the unfolded protein response. Cell. 1996;87: 391–404.

39. Plongthongkum N, Kullawong N, Panyim S, Tirasophon W. Ire1 regulated XBP1 mRNA splicing is essential for the unfolded protein response (UPR) in Drosophila melanogaster. Biochem Biophys Res Commun. 2007;354: 789–794. doi:10.1016/j.bbrc.2007.01.056

40. Ryoo HD, Domingos PM, Kang M-J, Steller H. Unfolded protein response in a Drosophila model for retinal degeneration. EMBO J. 2007;26: 242–252. doi:10.1038/sj.emboj.7601477

41. Shandala T, Woodcock JM, Ng Y, Biggs L, Skoulakis EMC, Brooks DA, et al. Drosophila 14-3-3ε has a crucial role in anti-microbial peptide secretion and innate immunity. J Cell Sci. 2011;124: 2165–2174. doi:10.1242/jcs.080598

42. Clemmons AW, Lindsay SA, Wasserman SA. An effector Peptide family required for Drosophila toll-mediated immunity. PLoS Pathog. 2015;11: e1004876. doi:10.1371/journal.ppat.1004876

43. Kenmoku H, Hori A, Kuraishi T, Kurata S. A novel mode of induction of the humoral innate immune response in Drosophila larvae. Dis Model Mech. The Company of Biologists Ltd; 2017;10: 271–281. doi:10.1242/dmm.027102

44. Fullerton MD, Hakimuddin F, Bonen A, Bakovic M. The development of a metabolic disease phenotype in CTP:phosphoethanolamine cytidylyltransferase-deficient mice. Journal of Biological Chemistry. American Society for Biochemistry and Molecular Biology; 2009;284: 25704–25713. doi:10.1074/jbc.M109.023846

45. Guo Y, Walther TC, Rao M, Stuurman N, Goshima G, Terayama K, et al. Functional genomic screen reveals genes involved in lipid-droplet formation and utilization. Nature. 2008;453: 657–661. doi:10.1038/nature06928

46. Leonardi R, Frank MW, Jackson PD, Rock CO, Jackowski S. Elimination of the CDP-ethanolamine pathway disrupts hepatic lipid homeostasis. Journal of Biological Chemistry. American Society for Biochemistry and Molecular Biology; 2009;284: 27077–27089. doi:10.1074/jbc.M109.031336

47. Lim HY, Wang W, Wessells RJ, Ocorr K, Bodmer R. Phospholipid homeostasis regulates lipid metabolism and cardiac function through SREBP signaling in Drosophila. Genes Dev. Cold Spring Harbor Lab; 2011;25: 189–200. doi:10.1101/gad.1992411

48. Walker AK, Jacobs RL, Watts JL, Rottiers V, Jiang K, Finnegan DM, et al. A conserved SREBP-1/phosphatidylcholine feedback circuit regulates lipogenesis in metazoans. Cell. 2011;147: 840–852. doi:10.1016/j.cell.2011.09.045

49. Harris CA, Haas JT, Streeper RS, Stone SJ, Kumari M, Yang K, et al. DGAT enzymes are required for triacylglycerol synthesis and lipid droplets in adipocytes. J Lipid Res. American Society for Biochemistry and Molecular Biology; 2011;52: 657–667. doi:10.1194/jlr.M013003

50. Yang C, Wang X, Wang J, Wang X, Chen W, Lu N, et al. Rewiring Neuronal Glycerolipid Metabolism Determines the Extent of Axon Regeneration. Neuron. 2020;105: 276–292.e5. doi:10.1016/j.neuron.2019.10.009

51. Harris TE, Huffman TA, Chi A, Shabanowitz J, Hunt DF, Kumar A, et al. Insulin controls subcellular localization and multisite phosphorylation of the phosphatidic acid phosphatase, lipin 1. J Biol Chem. American Society for Biochemistry and Molecular Biology; 2007;282: 277–286. doi:10.1074/jbc.M609537200

52. Cornell RB, Ridgway ND. CTP:phosphocholine cytidylyltransferase: Function, regulation, and structure of an amphitropic enzyme required for membrane biogenesis. Progress in Lipid Research. 2015;59: 147–171. doi:10.1016/j.plipres.2015.07.001

53. Infante JP. Rate-limiting steps in the cytidine pathway for the synthesis of phosphatidylcholine and phosphatidylethanolamine. Biochemical Journal. 1977;167: 847–849. doi:10.1042/bj1670847

54. Nyako M, Marks C, Sherma J, Reynolds ER. Tissue-specific and developmental effects of the easily shocked mutation on ethanolamine kinase activity and phospholipid composition in Drosophila melanogaster. Biochem Genet. Kluwer Academic Publishers-Plenum Publishers; 2001;39: 339–349. doi:10.1023/a:1012209030803

55. Weber U, Eroglu C, Mlodzik M. Phospholipid membrane composition affects EGF receptor and Notch signaling through effects on endocytosis during Drosophila development. Dev Cell. 2003;5: 559–570.

56. Meltzer S, Bagley JA, Perez GL, O’Brien CE, DeVault L, Guo Y, et al. Phospholipid Homeostasis Regulates Dendrite Morphogenesis in Drosophila Sensory Neurons. CellReports. 2017;21: 859–866. doi:10.1016/j.celrep.2017.09.089

57. Pavlidis P, Ramaswami M, Tanouye MA. The Drosophila easily shocked gene: a mutation in a phospholipid synthetic pathway causes seizure, neuronal failure, and paralysis. Cell. 1994;79: 23–33.

58. Gupta T, Schüpbach T. Cct1, a phosphatidylcholine biosynthesis enzyme, is required for Drosophila oogenesis and ovarian morphogenesis. Development. Oxford University Press for The Company of Biologists Limited; 2003;130: 6075–6087. doi:10.1242/dev.00817

59. Kirk SJ, Cliff JM, Thomas JA, Ward TH. Biogenesis of secretory organelles during B cell differentiation. J Leukoc Biol. Society for Leukocyte Biology; 2010;87: 245–255. doi:10.1189/jlb.1208774

60. Sriburi R, Bommiasamy H, Buldak GL, Robbins GR, Frank M, Jackowski S, et al. Coordinate regulation of phospholipid biosynthesis and secretory pathway gene expression in XBP-1(S)-induced endoplasmic reticulum biogenesis. J Biol Chem. 2007;282: 7024–7034. doi:10.1074/jbc.M609490200

61. Rosen DA, Seki SM, Fernández-Castañeda A, Beiter RM, Eccles JD, Woodfolk JA, et al. Modulation of the sigma-1 receptor-IRE1 pathway is beneficial in preclinical models of inflammation and sepsis. Sci Transl Med. 2019;11: eaau5266. doi:10.1126/scitranslmed.aau5266

62. Garsin DA. Ethanolamine utilization in bacterial pathogens: roles and regulation. Nat Rev Microbiol. 2010;8: 290–295. doi:10.1038/nrmicro2334

63. Romano KA, Martinez-Del Campo A, Kasahara K, Chittim CL, Vivas EI, Amador-Noguez D, et al. Metabolic, Epigenetic, and Transgenerational Effects of Gut Bacterial Choline Consumption. Cell Host and Microbe. 2017;22: 279–290.e7. doi:10.1016/j.chom.2017.07.021

64. Storelli G, Nam H-J, Simcox J, Villanueva CJ, Thummel CS. Drosophila HNF4 Directs a Switch in Lipid Metabolism that Supports the Transition to Adulthood. Dev Cell. 2019;48: 200–214.e6. doi:10.1016/j.devcel.2018.11.030

65. Hu X, Yagi Y, Tanji T, Zhou S, Ip YT. Multimerization and interaction of Toll and Spätzle in Drosophila. Proc Natl Acad Sci USA. National Acad Sciences; 2004;101: 9369–9374. doi:10.1073/pnas.0307062101

66. Yagi Y, Ip YT. Helicase89B is a Mot1p/BTAF1 homologue that mediates an antimicrobial response in Drosophila. EMBO Rep. EMBO Press; 2005;6: 1088–1094. doi:10.1038/sj.embor.7400542

67. Pascual A, Chaminade M, Preat T. Ethanolamine kinase controls neuroblast divisions in Drosophila mushroom bodies. Dev Biol. 2005;280: 177–186. doi:10.1016/j.ydbio.2005.01.017

68. Schmittgen TD, Livak KJ. Analyzing real-time PCR data by the comparative C(T) method. Nat Protoc. 2008;3: 1101–1108. doi:10.1038/nprot.2008.73

69. Bligh EG, Dyer WJ. A rapid method of total lipid extraction and purification. Can J Biochem Physiol. 1959;37: 911–917. doi:10.1139/o59-099

70. Serbulea V, Upchurch CM, Schappe MS, Voigt P, DeWeese DE, Desai BN, et al. Macrophage phenotype and bioenergetics are controlled by oxidized phospholipids identified in lean and obese adipose tissue. Proc Natl Acad Sci USA. 2018;115: E6254–E6263. doi:10.1073/pnas.1800544115

71. Tschanz SA, Burri PH, Weibel ER. A simple tool for stereological assessment of digital images: the STEPanizer. J Microsc. John Wiley & Sons, Ltd (10.1111); 2011;243: 47–59. doi:10.1111/j.1365-2818.2010.03481.x

72. Weibel ER, Kistler GS, Scherle WF. PRACTICAL STEREOLOGICAL METHODS FOR MORPHOMETRIC CYTOLOGY. J Cell Biol. Rockefeller University Press; 1966;30: 23–38. doi:10.1083/jcb.30.1.23

