## Supplemental Table 1 for "Innate immune signaling in *Drosophila* shifts anabolic lipid metabolism from triglyceride storage to phospholipid synthesis in an ER stress-dependent manner"

**S1 Table.** Genotypes of *Drosophila melanogaster* used in this study.

| Figure panels | Experiments, Genotype |
| --- | --- |
| ***Fig 1 and S1 Fig*** | ***Triglyceride measurements*** |
| 1A, 1B, S1A, S1B | w; (UAS-GFP or UAS-Toll$^{10b}$) / +; r4-GAL4 / + |
| 1C | w; +/+; (UAS-EGFP or UAS-Dif) / r4-GAL4 |
| 1D | w; +/+; (UAS-EGFP or UAS-Toll$^{10b}$, UAS-Dif$^{RNAi}$) / r4-GAL4 |
| ***Fig 2*** | ***Trehalose, glucose and glycogen measurements*** |
| 2A-2D | w; (UAS-GFP or UAS-Toll$^{10b}$) / +; r4-GAL4 / + |
| ***Fig 3*** | ***RT-qPCR and triglyceride measurements*** |
| 3B, 3C (left), 3D (left) | w; (UAS-GFP or UAS-Toll$^{10b}$) / +; r4-GAL4 / + |
| 3C (right), 3D (right) | w; +/+; (UAS-EGFP or UAS-Dif) / r4-GAL4 |
| 3E | w; (UAS-RFP / UAS-Toll$^{10b}$) / +; r4-GAL4 / (UAS-EGFP or UAS-LipinWT) |
| 3F | w; (UAS-RFP / UAS-Toll$^{10b}$) / +; r4-GAL4 / (UAS-EGFP or UAS-mdy$^{HA}$) |
| ***S2 Fig*** | ***RT-qPCR and triglyceride measurements*** |
| S2A | w; (UAS-GFP or UAS-Lipin$^{RNAi}$) / +; r4-GAL4 / + |
| S2B | w; (UAS-GFP or UAS-mdy$^{RNAi}$) / +; r4-GAL4 / + |
| S2C | w; (UAS-GFP or mdy$^{QX25}$) / +; r4-GAL4 / + |
| S2D | w; +/+; (UAS-EGFP or UAS-LipinWT) / r4-GAL4 |
| S2E | w; (UAS-GFP or P{EPgy2}mdy, CG13280) / +; r4-GAL4 / + |
| S2F | w; +/+; (UAS-EGFP or UAS-mdy$^{HA}$) / r4-GAL4 |
| S2G | w; (CyO, GFP or mdy$^{QX25}$) / +; (UAS-EGFP or UAS-mdy$^{HA}$) / r4-GAL4 |
| ***Fig 4 and S3 Fig*** | ***RT-qPCR and Western blot*** |
| 4B-4E, S3A, S3B | w; (UAS-GFP or UAS-Toll$^{10b}$) / +; r4-GAL4 / + |
| 4F, 4G | w; +/+; (UAS-EGFP or UAS-Dif) / r4-GAL4 |
| S3C | w; UAS-RFP / +; r4-GAL4 / UAS-EGFP (RNA-Seq: Suzawa et al., 2019) |
| ***Fig 4 and S3 Fig*** | ***Mass spectrometry and Thin Layer Chromatography*** |
| 4H, 4I, S3D, S3E | w; (UAS-GFP or UAS-Toll$^{10b}$) / +; r4-GAL4 / + |
| ***Fig 5 and S4 Fig*** | ***RT-qPCR and Western blot*** |
| 5A, 5C, 5D, S4E | w; Tub-GAL80ts / (UAS-RFP or UAS-Toll$^{10b}$); r4-GAL4 / (UAS-EGFP or UAS-Xbp1$^{RNAi}$) |
| 5B (left) | w; +/+; (UAS-EGFP or UAS-Dif) / r4-GAL4 |
| 5B (right) | w; +/+; (UAS-EGFP or UAS-Toll$^{10b}$, UAS-Dif$^{RNAi}$) / r4-GAL4 |
| S4A-S4D | w; (UAS-RFP or UAS-Toll$^{10b}$) / +; r4-GAL4 / (UAS-EGFP or UAS-SREBP1$^{RNAi}$) |
| ***Fig 6*** | ***RT-qPCR and electron microscopy*** |
| 6A-6D | w; (UAS-GFP or UAS-Toll$^{10b}$) / +; r4-GAL4 / + |
| ***Fig 7 and S5 Fig*** | ***RT-qPCR*** |
| 7A, S5A | w; (UAS-RFP or UAS-Toll$^{10b}$) / +; r4-GAL4 / UAS-EGFP (RNA-Seq: Suzawa et al., 2019) |
| 7B-7D, S5B | w; +/+ ; r4-GAL4, Drs$^{\Delta 7\text{-}17}$ / (UAS-EGFP or UAS-Toll$^{10b}$, Drs$^{\Delta 7\text{-}17}$) |
| 7B-7D, S5B | w; Bom$^{\Delta 55C}$ / Bom$^{\Delta 55C}$; r4-GAL4 / (UAS-EGFP or UAS-Toll$^{10b}$) |
| 7B-7D, S5B | w; Bom$^{\Delta 55C}$ / Bom$^{\Delta 55C}$; r4-GAL4, Drs$^{\Delta 7\text{-}17}$ / (UAS-EGFP or UAS-Toll$^{10b}$, Drs$^{\Delta 7\text{-}17}$) |
| 7C, 7D | w; (UAS-GFP or UAS-Toll$^{10b}$) / +; r4-GAL4 / + |
