## Supplemental Table 2 for "Innate immune signaling in *Drosophila* shifts anabolic lipid metabolism from triglyceride storage to phospholipid synthesis in an ER stress-dependent manner"

**S2 Table.** Sequences of oligonucleotides used in this study.

| **qRT-PCR primers, sequences listed 5' → 3'** | | |
|---|---|---|
| **gene** | **Forward primer** | **Reverse primer** |
| Rp49 | CGCTTCAAGGGACAGTATCTG | AAACGCGGTTCTGCATGA |
| ATPCL | AAGGACATCCTGAACCGCCAT | GGCATTTCCGATCTTCTGGTC |
| ACC | CGCTATGGTTACCTGCCGTA | CGCTATGGTTACCTGCCGTA |
| FASN | GTGTTGGCCAACATTGACTAC | CTCAACAGTGTGTACTTCCAC |
| Lipin | GCGAGGTGATTGAGAAGAAG | AATGTTTGGGTCAGCTCGTG |
| midway | ACTGCTCTGCATTGGAGGTC | ATGTCCTTCGCCTTCGTTGT |
| eas | ACCAAACCCATGCCGATGAT | GGCGGCCGATAGGTAGAAAC |
| Pect | CAACGATGGAGGCAAATGGC | GGCATGTCCGAAGTGTACCA |
| CG7149 | GAGGCTCATTTGAGGGGCTT | AGGGTGCATCACATAGACGC |
| CG2201 | CACACAAACGTCAGCGATGC | ATTCGCAGAAGGACCTCACG |
| Pcyt1 | TTACCTCAACGGCGTCAAGC | AAATTGTCGCAAAATCGGAGGC |
| bbc | GCGGTTCGGCAGAATTGATG | GACACTGGTGCCCTTGGTG |
| Pcyt2 | AGCAATGGACAGAATGCCGA | GCAGGCTGGCAAATGCTAAAA |
| CG33116 | ACGCAGGACCAGATCAATGG | CCAGCGCGGAAAGAACTTAAC |
| SREBP | CCTCTGCGATGAGTCGAGTG | GCCAATCGCAGGTAAGCAAC |
| Xbp1u | TCTGCAGCATCCAAAGCTGAC | GCATGTCTTGTAGAGTAGGC |
| Xbp1s | CTTGGATCTGCCGCAGGGTAT | GCATGTCTTGTAGAGTAGGC |
| BiP | ATATTACTGGCCGTCGTGGC | ACCAACGCAGGAATACGTGG |
| Pdi | GACGGTGGACAACTTCAAGC | CTTGATGGGCGACTCCTTCTC |
| Edem1 | GCGGATATGCAACGATTCGC | GACCCATCGTTGTGCAGGAA |
| Drs | AGTACTTGTTCGCCCTCTTCG | GGTCTCGTTGTCCCAGACG |
| BomS2 | TGGCCAACGCTGTTCCC | CCTACTTTCCACCGTGCACAT |
| IM4 | CAAGCCAACCAACAACCACC | ATGAGCACGGTTCCAGGTTG |
| IM14 | TCTGCGGCTTTTTCTTCGCT | TTTGAATCAACGTGTGTCCGC |
| **Generation of UAS-HA.mdy** | | |
| ENTR HA mdy-F | CACCATGTACCCATACGACGTCCCAGACTACGCTACCACCAATAAGGATCCCCAAGATAAG | |
| mdy-R | CTAACTACTGTAGTCGGTGCCGTTGA | |
